# Individualised niches in a variable environment – Consequences for environmental change responses

**DOI:** 10.1101/2025.03.24.644970

**Authors:** Anastasiia Enne, Vishnu Venugopal, Peter Nabutanyi, Meike J. Wittmann

**Affiliations:** Bielefeld University

## Abstract

1. Intraspecific trait variation (ITV) can be important for population performance in a variable and changing environment because individuals with different traits have different fitness responses. Furthermore, there are three mechanisms via which individuals can interact with their environment to potentially improve fitness: niche conformance, niche construction, and niche choice (NC^3^). These processes become increasingly important in the presence of environmental change, but there is still no mathematical modelling framework that would unite the effects of ITV and the NC^3^ mechanisms.
2. In this paper, we build a general model incorporating ITV and two of the NC^3^ mechanisms (niche conformance and construction, NC^2^) to investigate how they affect populations in a changing and variable environment via non-linear averaging. We quantify the effects of NC^2^ and ITV on average individual fitness using an analytical Taylor approximation and a sampling approach.
3. Our method allows us to answer the question of what would have happened if individuals in the study system did not have ITV or did not perform NC^2^ mechanisms. The answer to this question depends on the curvature of the fitness function and can be estimated via the Taylor approximation.
4. We apply the method to two case studies: great tits adjusting their laying date to yearly changes in vegetation green-up, and a host-parasite system in which the parasite changes its environment by immunodepression of the host. In the great tits, we found a slight negative effect of ITV and a slight positive effect of niche conformance on the population fitness. In the host-parasite system, we found ITV to have no effect without niche construction, but with niche construction ITV decreased virulence. Also, niche construction had a strong negative effect on virulence.
5. Our extension of non-linear averaging theory, combining intraspecific and environmental variation, niche conformance, and niche construction, allowed us to assess average population performance with those mechanisms at play. However, how well one can estimate such performance depends on the type of data available. This framework can be extended further to niche choice and evolution, therefore including all processes that can change the match between individuals and their environment.

## 1 Introduction

Better understanding how populations respond to variable and systematically changing environmental conditions is key for predicting and protecting biodiversity under global change. Most previous models have made such predictions by using a single fitness function as a measure of the species’ ecological niche, e.g. a thermal performance curve or temperature niche limits for the whole species (Bernhardt *et al*., 2018; Meyer *et al*., 2022; Khelifa *et al*., 2019). However, in most species, there is considerable intraspecific trait variation (ITV) due to ontogeny, genetic variation, epigenetic variation and plastic responses to environmental variation (Moran *et al*., 2016; Westerband *et al*., 2021). Individuals with different trait values can have different individualized niches, that is, different fitness functions specifying the sets of environmental conditions under which the individuals are able to thrive (Layman *et al*., 2015; Takola & Schielzeth, 2022). Thus, for accurate predictions of population performance in a variable (in space and over time) environment, ITV needs to be taken into account, as has already been shown in some ecological niche models (Razgour *et al*., 2019; Kearney *et al*., 2009; Mathewson *et al*., 2017).

Furthermore, an individual’s traits and the environmental or micro-environmental conditions it experiences are not necessarily fixed. Recent studies have distinguished three types of individual-environment interaction via which an individual can improve the fit between its trait and its (micro-)environment within its own lifetime (Trappes *et al*., 2022; Edelaar & Bolnick, 2019; Aaby & Ramsey, 2022). These so-called NC^3^ processes (Trappes *et al*., 2022) are 1. niche choice where individuals select a (micro-)environment that matches their traits better, 2. niche conformance (phenotypic plasticity) where individuals change their phenotype in response to the given (micro-)environment, and 3. niche construction when individuals change the properties of their (micro-)environment. Note that with the individual’s niche defined as its fitness function, only niche conformance changes the niche, while niche construction and choice change just the environment experienced (Trappes *et al*., 2022). In many species, individuals use two or more of the NC^3^ processes. For example, decomposing fungi do not only modify the soil environment via plant matter decomposition (niche construction, Odling-Smee *et al*., 2003), but also adjust their extracellular enzyme activity in response to changes in temperature and humidity (niche conformance, Alster *et al*., 2021). Female meerkats adjust their food intake to their social niche while subordinate (niche conformance, as reviewed in Komdeur & Ma, 2021), but shape their social environment later in life as dominant females (niche construction, Young *et al*., 2006). In this paper we focus on the consequences of ITV, as well as niche construction and niche conformance (abbreviated as NC^2^) for population performance in a variable environment.

Over the last years, empirical evidence has been accumulating that ITV can have strong and far-reaching ecological consequences (Bolnick *et al*., 2011; Violle *et al*., 2012; Des Roches *et al*., 2018; Cope *et al*., 2022; Giménez & Jenkins, 2024). Across various taxa (plants, animals, bacteria), ITV increases population performance in the face of various types of environmental change, e.g. temperature stress, invasive species, or parasites (Forsman & Wennersten, 2015). For many mammalian populations, reduced variation in developmental, morphological, reproductive, and ecological traits increases extinction risk (Gonzalez-Suarez & Revilla, 2013). ITV also plays an important role in species coexistence when affecting species interaction parameters. It increases network stability (Moran *et al*., 2016) and improves response of a network to recovery measures (Baruah & Wittmann, 2024). However, networks with high ITV may also sometimes collapse more abruptly (Baruah, 2022). High genetic diversity can also increase colonization or establishment rates, and reduce the rate of extinction, by reducing resource competition or increasing the probability of the presence of the more successful individuals (Moran *et al*., 2016; Forsman & Wennersten, 2015).

Among several mechanisms mediating the ecological consequences of ITV (Bolnick *et al*., 2011), one very general mechanism operating irrespective of the trait’s mechanistic or genetic basis is non-linear averaging. Suppose there is trait variation among individuals (e.g. in body size) and we are interested in the population average of a demographic parameter such as the number of offspring. If at the individual level, the demographic parameter is a non-linear function of the individual trait (e.g. the number of offspring as a function of body size) or a non-linear multivariate function of multiple traits, then the population average of the function (e.g. number of offspring) is in general not equal to the function of the population average trait value(s). The direction of the effect can be predicted based on Jensen’s inequality and the curvature (second derivative) of the function of interest in the region around the species’ trait mean (Jensen, 1906). If the function is concave down (the second derivative is negative), then the population average of that function is smaller than the function of the population average trait value. Vice versa if the function is concave up (the second derivative is positive), then the population average of that function is greater than the function of the population average trait value. Previous theory has shown that that via non-linear averaging, ITV can have strong effects on interspecific interactions, for example between competitors (Hart *et al*., 2016; Uriarte & Menge, 2018) or between predators and their prey (Okuyama, 2008; Gibert & Brassil, 2014; van Benthem *et al*., 2024). We also know that non-linear averaging plays an important role when it comes to environmental variability. Past studies have found effects of temperature variability on population growth (Bernhardt *et al*., 2018; Pickett *et al*., 2015), and effects of environmental variability on species coexistence (Gravel *et al*., 2011). Jensen’s inequality also predicts negative effects of temporal variation in light intensity (irradiance) for a plant’s ability to photosynthesise (Ruel *et al*., 1999), and reduced performance for individuals in an environment with more variable temperature (Denny, 2019).

In addition, previous theory suggests that niche construction and niche conformance can often buffer the negative consequences of environmental change (Moran *et al*., 2016; Steinmetz *et al*., 2020; Coninx, 2023). Niche conformance, even if it may slow the genetic response to environmental change, can allow populations to cope with faster rates of environmental change (Chevin *et al*., 2010; Forsman, 2014). The role of a plastic response becomes even more important in human-altered environments, where changes happen quicker (Chevin *et al*., 2012). Similarly, niche construction may allow species to persist under harsher conditions than they would be able to sustain without niche construction (Kylafis & Loreau, 2008), sometimes affecting whole ecosystems (Brathen *et al*., 2018). This makes accounting for niche conformance (Chevin *et al*., 2012) and niche construction in ecological modelling highly important. However, niche conformance and construction are not always beneficial. For example, niche conformance can have a fitness disadvantage when environmental cues are not reliable (Reed *et al*., 2010), such as when *Physella virgata* snails flee in the presence of the sunfish even if it poses no risk, and produce a more round shell that leads to slower growth of individuals (Miner *et al*., 2005). As another example, green sea turtles *Chelonia mydas* experience dramatic skews in sex ratios due to climate change coupled with temperature-dependent sex determination (Hill *et al*., 2024).

Given these mixed effects of ITV, niche conformance and construction, finding general rules for the effects of ITV combined with the NC^2^ mechanisms is challenging. A few important aspects are missing so far in theories for the effect of ITV and NC^2^ under environmental change. First, most models assume that all individuals at a given time experience the same environment, but in reality each individual has its own micro-environment. We thus need to take a two-dimensional (micro-environment vs. individual trait) perspective. Second, non-linear averaging using a Taylor approximation to find the population average of the non-linear fitness function is a potentially powerful tool to study the role of ITV in response to environmental change, but, to our knowledge, has only been used to model the effects of environmental variation without ITV (see e.g. Chesson, 2009; Bernhardt *et al*., 2018). Furthermore, non-linear averaging theory so far has not taken into account niche construction or niche conformance.

To help fill these knowledge gaps, our goal is to build a general model that incorporates

- effects of individual trait variation (ITV),
- effects of niche conformance and construction (NC^2^ mechanisms)

to predict consequences for average population fitness in a world where there is variation among micro-environments experienced by individuals and also systematic change in the average environmental conditions. The novelty of our approach is in bringing NC^2^ mechanisms and Taylor approximations together with two-dimensional trait-environment or trait-trait space (meaning, that a trait of an individual of one species can be an environment for an individual of another species, e.g. in a host-parasite system). The desired outcome is a 2D non-linear averaging theory of environmental change responses, that helps us gain more understanding of the interplay between these mechanisms.

We are using a simple model with individual fitness as a function of trait and micro-environment. We then can compute average population fitness under changing environmental conditions using the second-order Taylor approximation and use this to assess average population fitness under the given conditions. We illustrate our modelling framework with two case studies. For a study system of great tits in a changing environment we estimate the individual fitness function and the strength of the niche conformance mechanism to find out how important niche conformance is and how ITV affects individual fitness. For a host-parasite system, we estimate the individual fitness function (measured as virulence) and the strength of the niche construction mechanism of the parasite to find out how important niche construction is and how ITV affects virulence.

## 2 Models and methods

Here we introduce our mathematical framework to model effects of ITV and the NC^2^ mechanisms for population fitness (Fig. 1). As a brief overview, we model individual fitness as a function of individual trait and micro-environment (figure 1A) and we consider NC^2^ mechanisms as functions transforming individual trait (niche conformance) and environment (niche construction)(Fig. 1B). To find the population average fitness, we use an analytical approximation that takes into account the fitness function (and its curvature), and the niche conformance/construction functions. This integrates our way of modelling NC^2^ mechanisms with a second-order Taylor approximation (Gravel *et al*., 2011), which allows us to study the effects of ITV and NC^2^, and their interactions together in one modelling framework. To complement this approach, we use a stochastic sampling method as another way of quantifying effects of NC^2^.

**Figure 1.**
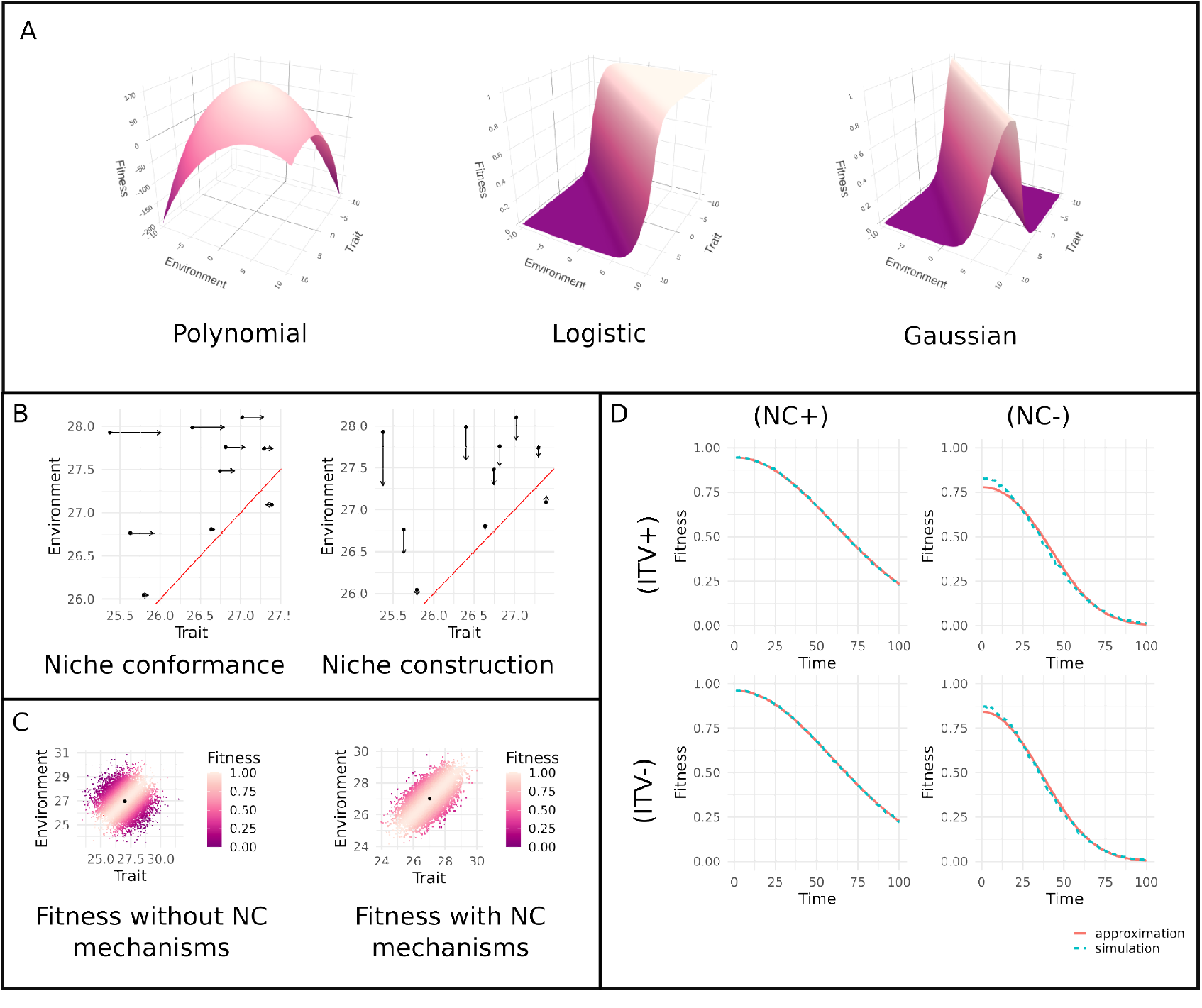
Key components of the modelling approach. **A)** Examples of fitness functions. **B)** How niche conformance and construction affect individuals in a trait-environment space. The red line is the perfect match between trait and environment, black dots are individuals, arrows show the direction and distance at which niche mechanisms “move” individuals in a trait-environment space. The niche conformance function and niche construction function are assumed to be *z*′ = *g*(*z, e*) = *z* + 0.25(*e* − *z*) and *e*′ = *h*(*z, e*) = *e* + 0.25(*z*− *e*), respectively. **C)** How such niche conformance and construction affect population fitness. Each point is an individual and the colour of the points shows the value of individual fitness. **D)** Effects of NC^2^ and ITV in a changing environment. We take a population of 1000 individuals over 100 time steps. Over these time steps the average micro-environment value increases linearly. The first row corresponds to the presence of ITV; the second row corresponds to the absence of ITV, the first column corresponds to the presence of niche conformance and construction (NC^2^); the second column corresponds to the absence of *NC*^2^. The solid red line is the analytical approximation of the average population fitness. The dashed blue line is average fitness from sampling 1000 individuals and calculating the sample mean. The script to obtain the figure can be found in the code supplementary.

For this framework, we consider only one trait and one environmental parameter. We also assume in this simple model that the trait distribution before the action of niche mechanisms stays constant across generations, i.e. there is no heritable component. This is the first step towards developing a more detailed model with heritable variation, and can be realistic, for example, for cases with random developmental variation and ontogenetic variation. We consider the fitness function to be at least twice differentiable. For the analytical approximation, we need to know averages and standard deviations of individual trait and environmental values, but we are not bound to any specific probability distribution. The approximation part of the framework is sensitive to variance in trait and environment, but as long as it is not too large compared to the scale at which the curvature of the fitness function changes, it is an adequate way to find average fitness.

Suppose that each individual *i* has a trait *z*_*i*_ drawn from a distribution with mean *z* and standard deviation *σ*_*z*_, and lives in a micro-environment *e*_*i*_ drawn from a distribution with mean *e* and standard deviation *σ*_*e*_. Then the fitness of this individual (e.g, number of descendants) is denoted as *f* (*z*_*i*_, *e*_*i*_).

When an individual performs niche conformance, it changes its trait based on its given trait and micro-environment (figure 1B). Many studies consider the reaction norm or niche conformance function as a linear dependence between trait and environment (e.g., Westneat, 2024; Cole *et al*., 2015). We can write this relationship as 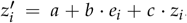. Geometrically, this means that the function *g* is a linear transformation of trait-environment space (*z, e*). Transformation happens only along the trait axis, since only the trait is changing (see figure 1B). More generally, the conformance function can be any twice differentiable function *g*(*z*_*i*_, *e*_*i*_) (e.g., polynomial in Morrissey & Liefting, 2016). For each individual, we can calculate a new 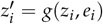 based on a given trait *z*_*i*_ and micro-environment *e*_*i*_. This new trait will then be substituted into the fitness function 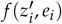 and change the individual’s fitness (see figure 1C).

The same approach can be used for the niche construction function, only this time the environment is changing: 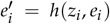.Again, this new micro-environmental value is plugged into the fitness function 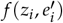 to change the individual’s fitness in the presence of niche construction. Similar to niche conformance, function *h*(*z*_*i*_, *e*_*i*_) should be twice differentiable. Some studies use linear equations to describe niche construction (e.g., Brock *et al*., 2016; Odling-Smee *et al*., 2003), while others opt for a non-linear form (Kylafis & Loreau, 2008). Here, in the case of a linear niche construction function *e*′ = *h*(*z, e*) = *a* + *b* · *z*_*i*_ + *c* · *e*_*i*_, it will move individuals along the environment axis in a trait-environment space (*z, e*) (see figure 1B).

In our framework, when we assume that individuals perform both NC^2^ mechanisms, we assume that they perform them simultaneously. That is, trait and environment change at the same time, resulting in new trait *z*′ = *g*(*z, e*) and new micro-environment *e*′ = *h*(*z, e*), which then affect the fitness. This may result in better trait-environment match as individuals move along both axes in trait-environment space (*z, e*) (see figure 1C).

For dynamical models (i.e., models where we are looking at how population size changes over time) or more generally to assess the current performance of the population, we usually want to know population average fitness. In our paper we quantify average fitness with two different approaches: a sampling method and an analytical approximation. The sampling method requires sampling a large number of individuals and micro-environments, applying the niche conformance and construction functions, calculating fitness for each individual, and then computing an average of all fitness values. A drawback of this approach is that we do not get an analytical expression of the direction and magnitude of the separate fitness effects of ITV, niche conformance and niche construction.

The analytical approximation estimates average fitness based on the shape of fitness function. First, let us consider a second-order Taylor approximation of *f* around the mean trait and mean environmental value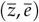:

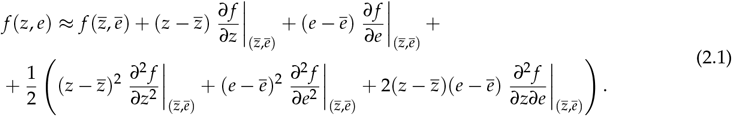

Taking the expectation of this expression allows us to find the expected fitness with ITV and micro-environmental variation, but without NC^2^ mechanisms:

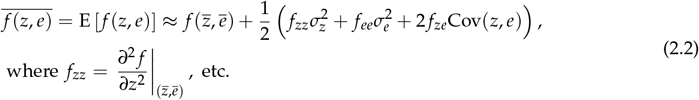

Throughout this paper, we will be assuming that Cov(*z, e*) = 0. This means, that trait and environment are linearly independent. In particular, there is no tendency among individuals to end up in the environment they prefer. One can drop this assumption to include the third NC mechanism – niche choice-into the framework. Other works have used similar approaches to find, for example, long-term average growth rate (Gravel *et al*., 2011), integrated physiological performance in a variable environment (Koussoroplis *et al*., 2017), or average interaction parameters in a species interaction (van Benthem *et al*., 2024). Here we show how NC^2^ mechanisms can be included into the analytical approximation.

To take into account the effects of the NC^2^ mechanisms, we need to approximate not a simple function *f* (*z, e*), but a more complex one *F*(*z, e*) = *f* (*g*(*z, e*), *h*(*z, e*)). We can find the second-order Taylor approximation for such a function (see equation S1.1 and section S1.1 for the detailed derivation of this formula). Based on this, we find the expectation

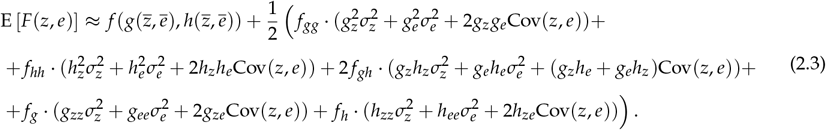

Note that 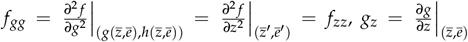, etc. To find derivatives of *f* with respect to *g* and/or *h* in our case we can just find derivatives of *f* with respect to *z* and/or *e* and then substitute 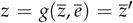 and 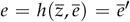 (*z*′ and *e*′ denote the trait and environment after NC^2^ mechanisms respectively; note that in case of nonlinear conformance or construction functions, *z*′ and *e*′ are not necessarily equal to the mean trait after conformance and the mean environment after construction). Using this, we can rewrite (2.3) as

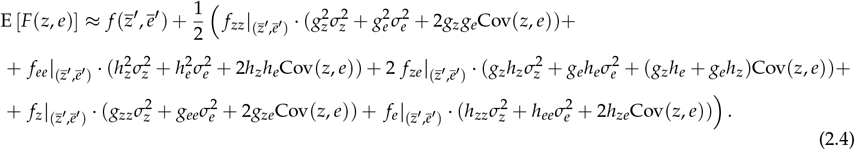

We provide a small example illustrating the approximation approach in figure 1D. We take a population of 1000 individuals and assume they have temperature tolerance as trait *z*, temperature as micro-environment *e*, and a Gaussian fitness function 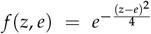. We take 100 time steps and change the average micro-environment from *ē* = 27 to *ē* = 32 over them. The population average of the trait stays constant at 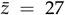.For scenarios with NC^2^ mechanisms, we assume that the niche conformance function is *z*′ = *g*(*z, e*) = *z* + 0.25(*e* − *z*) and the niche construction function is *e*′ = *h*(*z, e*) = *e* + 0.25(*z* − *e*). For scenarios without NC^2^ mechanisms, these functions are *z*′ = *g*(*z, e*) = *z* and *e*′ = *h*(*z, e*) = *e*, respectively. For scenarios with ITV, we assume that the trait has standard deviation 0.5, and for scenarios without ITV we set the standard deviation to 0. The environmental standard deviation stays constant and equal to 0.8. For each time step, we calculate average fitness based on the sampling method (via sampling of individual traits and micro-environments, letting them perform the NC^2^ mechanisms, calculating the individual fitness, and then averaging it over 1000 individuals), and the analytical approximation (by substituting derivatives of fitness and NC^2^ functions along with mean and standard deviation of the trait and environment into the equation 2.4). In this toy model, NC^2^ mechanisms help the population to preserve higher population fitness for longer. ITV does not benefit the population at the very beginning, when it is still well adapted to the environment, but further away from this point it gives a slight fitness advantage. The approximation basically coincides with the results from the sampling approach with NC^2^ mechanisms, with slight discrepancies without, because the deviations from the average trait and environment are larger in that case.

Our approach can be applied to empirical data from the field or laboratory to investigate the role of ITV, environmental variation, niche conformance, and niche construction for average population fitness. We here briefly describe the general approach (outlined in figure 2) and in the next section illustrate it in detail with two specific case studies. The first step is to obtain data on trait, environment and a fitness proxy for a large number of individuals. Ideally, we would also have trait values before niche conformance and environmental values before niche construction for all individuals, but some workarounds are possible when this information is not available or only partially available (see case studies below). The second step is to fit fitness and niche conformance and/or construction functions to this data (see figure 2B,C). Below, we will use GLMM in one case study and non-linear least squares in the other to show how our approach can flexibly accommodate different model fitting methods. Also splines (Wetzel *et al*., 2016) or a tensor product smooth (Wood, 2006; van Benthem *et al*., 2024) would be possible. It is important that the fitting approach is flexible enough to allow e.g. the fitness function to take various shapes with different curvatures, as informed by the data. If there are multiple candidate models, one additionally has to perform model selection. The third step is to use the estimated functions to derive average population fitness using either the analytical approximation or the sampling approach. To find out how ITV, environmental variation, and the NC^2^ mechanisms affect average fitness, we have to investigate hypothetical scenarios in which one or more of these are deactivated. One way to account for the hypothetical case of absence of ITV is to use the analytical approximation in equation 2.4 and substitute 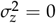.Another way would be to take a sample of individuals with no trait variation and calculate average fitness over this sample. The same principles can be applied when we want to compare presence and absence of NC^2^ mechanisms. Using the niche conformance and niche construction functions, we calculate how individual traits or micro-environments would look like without NC^2^ mechanisms. In the sampling method, we then sample individuals with such trait or micro-environment distribution. In the analytical approximation, we substitute *g*(*z, e*) = *z* or *h*(*z, e*) = *e* into equation 2.4. Finally, one can interpret the results in the context of the biological system and assess whether ITV and the NC^2^ mechanisms have positive or negative effects on average

**Figure 2.**
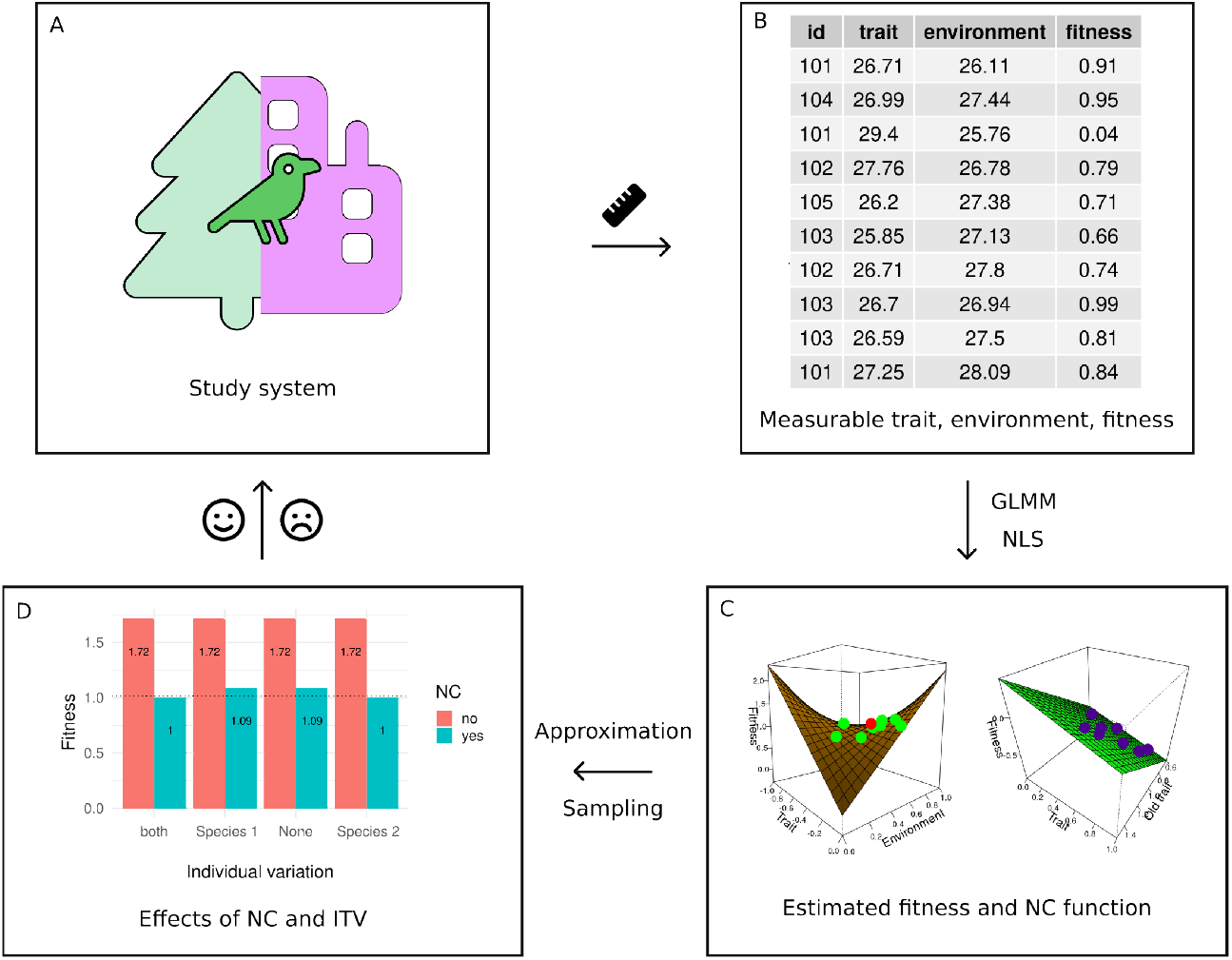
The workflow for applying our model to an empirical study system. We start with the study system **(A)**, that has data on trait, environment, and fitness, possibly having some individuals appearing multiple times in the dataset **(B)**. Using GLMM, non-linear least squares, or similar methods, we estimate fitness and NC^2^ functions **(C)**, and then consider effects of NC^2^ and ITV **(D)** via approximation or sampling methods to assess how well the population is doing **(A)**. In C and D, figures from case study 2 are inserted for illustrative purposes. fitness.

## 3 Case studies

### 3.1 Niche conformance of great tit laying dates

We apply our method (see figure 2) to a case of phenological mismatch caused by climate change in great tits (*Parus major*). Individuals must properly time their reproduction in a seasonal environment to provide their offspring with enough food during crucial developmental stages of their lives (Visser *et al*., 1998). Great tit nestlings reach peak food demand approximately 9-10 days after hatching (Reed *et al*., 2013). Thus parents need to match this peak demand to the period when the main food source – caterpillars – is at its highest abundance. The caterpillar peak can be measured as caterpillar half-fall date (date on which half of the caterpillars have dropped from the trees; Van Noordwijk *et al*., 1995).

Caterpillar half-fall date has been shown to positively correlate (Cole *et al*., 2015) to vegetation greenup date (date when EVI2, the two-band enhanced vegetation index, first crossed 15% of the segment EVI2 amplitude; Friedl *et al*., 2022), which we use as our environmental variable. Climate change can affect individual fitness by increasing the mismatch in time between hatching chicks and emergence of caterpillars (Reed *et al*., 2013). The main mechanism that allows great tits to adjust to the changing climate is phenotypic plasticity, i.e. niche conformance (Charmantier *et al*., 2008; Vedder *et al*., 2013; Cole *et al*., 2015). We explore how niche conformance and ITV affect population fitness by estimating fitness and niche conformance functions from the data set published in Cao *et al*. (2019) and then quantifying average individual fitness with and without ITV and niche conformance, either via the sampling approach or using the analytical approximation (equation 2.4). We provide more details of this analysis in section S2.

To model effects of niche conformance on population fitness, we estimate our model variables (egg laying date as individual trait *z*, green-up date as environmental variable *e*, and number of fledglings as fitness) using two data sets. First, we use the data set published in Cao *et al*. (2019) for the great tit population in the Hoge Veluwe National Park in the Netherlands. Each of the 5892 records has information about breeding mother identity, year of breeding, laying date, number of fledglings, and brood type (first brood, replacement brood, second brood). We narrowed down the data by focusing on the first brood only and we removed all the records with breeding year earlier than 2001, because for estimating green-up date we used NASA satellite data (Friedl *et al*., 2022), which does not exist before 2001. Second, we add environmental data to this data set. For each year, we take an average green-up date from a rectangle drawn on top of the study site (52°02′ – 52°07′N, 5°51′ – 5°32′E; Cao *et al*., 2019). Our trait – egg laying date – has an average value of 19.84079 and a standard deviation of 7.642885. Across all years, our environmental variable – green-up date – has an average value of −12.44094 and a standard deviation of 7.613845. All dates are in days after March, 31 (with negative dates being in March).

To estimate the fitness function, we fitted a zero-inflated Poisson generalized linear mixed model (GLMM) with logit link following the works of Chevin *et al*. (2015) and Cole *et al*. (2015). Our response variable was the number of fledglings and our fixed effects were *z*^2^, *e*^2^, *z* · *e, z, e* and the random effect was breeding mother’s ID. In order to capture fitness functions that are of Gaussian or similar shape we included trait and environment terms in the equation up to a second order. We used automated model selection on different combinations of trait and environment terms with the R package MuMIn (Barton, 2023), ranking the models according to their Akaike Information Criteria (AIC). We also checked other model types (Poisson, negative binomial, truncated Poisson), but they all had higher AIC. The general outcome of the model fitted is of the form

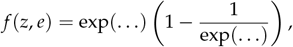

where … stands for fixed effects (in the first exponent for the count part, and in the second one for the zero-inflated part; the full equation is provided in section S2.6). The zero-inflated Poisson model assumes that in addition the Poisson-distributed number of fledglings, which can produce zero fledglings (count part), there are other processes that can result in zero fledglings, but are not tied directly to the mother’s quality, e.g. predation. Thus, the zero-inflated part can be interpreted as the probability that this second group of processes will not result in zero fledglings. The best model according to our selection criteria had environment (*e*), trait (*z*), *z*^2^, and *z* · *e* terms in the first exponent and all possible terms up to the second order in the second exponent, and no random effects (more details are provided in section S2.6).

Cole *et al*. (2015) find a linear relationship between laying date and green-up date, estimating the slope of the reaction norm. We find this relationship between trait and environment by fitting a simple linear regression with laying date as response variable (assuming that the observed laying date is the trait after niche conformance *z*′) and *e* as the fixed effect and get it in the form

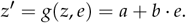

We do not know trait values prior to niche conformance (i.e. how the trait values would look like had individuals not performed niche conformance), but we can assume that if we remove part of the trait that is explained by the environment, then what is left (*a*) is the inherent individual trait without niche conformance. Then trait values prior to niche conformance (*z*) are

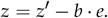

See section S2.5 for more details.

Given the estimated niche conformance and fitness functions, we can now use our analytical approximation and the sampling approach to find population-level average fitness. The expression for average fitness (number of fledglings) with niche conformance reduces to (compare to equation 2.4):

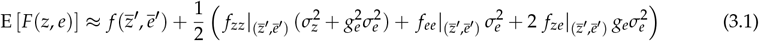

This happens because we do not consider niche construction (hence, *e*′ = *h*(*z, e*) = *e*), and the niche conformance function is linear with respect to *z* and *e*. Here, 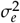 is between-year variance of the environment (since our framework does not have generational or yearly structure we can use the available source of variation – between year variation). We used a bootstrapping approach to quantify uncertainty in our model estimates. For that we generated 100 bootstrap samples from the dataset and repeated the procedure of fitting the zero-inflated Poisson GLMM for the fitness function and a linear model for the niche conformance function. The results of the bootstrapping method are shown as error bars for the approximation and sampling methods in figure 3, and more details on the method are provided in section S2.6.

**Figure 3.**
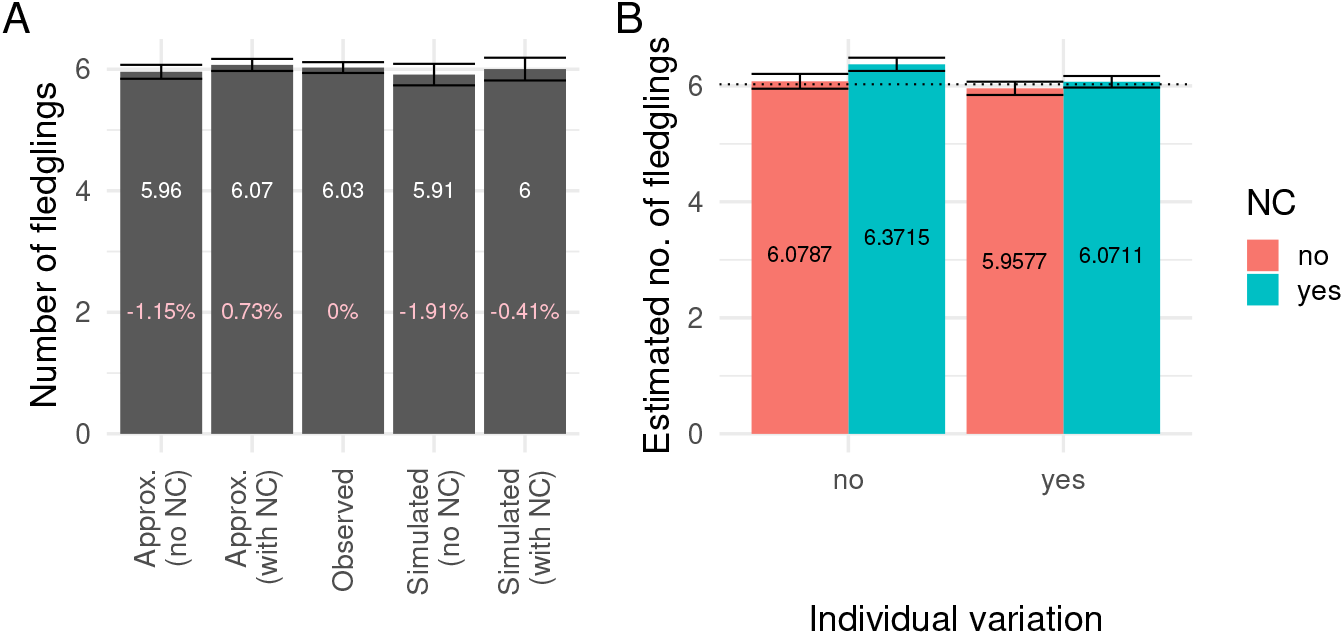
(**A**) Effect of niche conformance on average fitness. Bars represent (from left to right) approximation of average fitness without niche conformance, approximation of average fitness with niche conformance, observed average fitness, simulated average fitness without niche conformance, simulated average fitness with niche conformance. White numbers on the bars are values of average fitness for each bar, pink numbers on the bars are difference between bar value and observed average fitness in percentages of the observed average fitness. Error bars represent uncertainty due to bootstrapping procedure as standard deviation (for approximation and simulated average) and standard error of the mean (for observed average). (**B**) Effect of ITV and niche conformance on individual fitness according to analytical approximation. Bars represent approximated average fitness. The two leftmost bars represent the case without ITV, the two rightmost bars represent the case with ITV. The two red bars represent the case without niche conformance, the two green bars represent the case with niche conformance. Numbers on the bars are values of approximated average fitness for each bar. Error bars represent uncertainty due to bootstrapping procedure as standard deviation. The dashed horizontal line is the observed average fitness.

We calculated approximated average population fitness with and without niche conformance. We also calculated average population fitness using the sampling method (simulated in figure 3A): drawing random trait values for all individuals from the normal distribution (with mean and standard deviation equal to those of the dataset) and calculating actual average fitness for 100 replicates. Absence of niche conformance slightly decreases average population fitness. The approximation estimates well the average fitness for the observed data as well as the sampling approach, although error bars for both scenarios in each approach overlap, thus making the difference between scenarios less significant, and making it hard to give a very clear conclusions about the effects of NC and ITV.

According to the analytical approximation, average individual fitness would be higher in the absence of ITV for the estimated fitness function (figure 3B). Without niche conformance our analytical approximation simplifies to

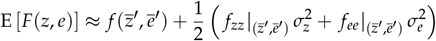

where *f*_*zz*_ is negative. Hence the curvature of our fitness function (see figure S2.9) leads to negative effects of ITV on average fitness.

We also performed the same analysis for each year separately (see reffig:hist1yearly and S2.12). For individual years, both methods worked less predictably, since the fitness function was fitted for the whole dataset. The difference from observed average fitness was both in negative and positive directions, effects of niche conformance were also both negative and positive, effects of ITV remained mostly negative.

### 3.2 Niche construction of the host micro-environment by a parasite

Host-parasite systems provide several examples of niche construction like parasites manipulating their host’s behaviour (Hindsbo, 1972; Holmes & Bethel, 1972) or other host phenotypes (Gopko & Mikheev, 2019). In these examples, the individual host can be seen as the parasite’s micro-environment and any parasite modification of this environment can be viewed as niche construction.

We base our case study on an empirical study (Cornet *et al*., 2009b) on the effects of variation in parasite infectivity and variation in host immunity on the virulence in the *Gammarus pulex* (amphipod)*Pomphorhynchus laevis* (acanthocephalan) system where the acanthocephalan is the parasite and the amphipod is the host (in this case an intermediate host, that is then consumed by fish as definitive host). Virulence, defined as the host mortality, has a negative consequence on the parasite’s fitness (Leggett *et al*., 2013), and we consider it as a fitness measure of the parasite. Being an emergent property of the host-parasite interaction, in this case study, we consider it as a function of the parasite trait infectivity and the host trait immunity. Virulence (*V*), immunity of host (*e*) and infectivity of the parasite (*z*) thus correspond to fitness, environment and trait variables in section 2.

Infection by *P. laevis* is shown to weaken the immune defence of the host by impairing the effectivity of prophenoloxidase (proPO) and circulating haemocytes (proxies for host immune system function) (Cornet *et al*., 2009b,a). Upon infection of the host by the parasite, virulence was found to be positively correlated to the immune expression of the host (proPO). In other words, the higher the level of immunodepression by the parasite on the host, the higher was the host survival. Immunopathology, the overreacting immune system of the host triggered by the infection harming the host tissues, was thought to be a cause for this observation.

To estimate the fitness function, we fitted various candidate polynomial models with virulence (*V*) as the response variable and infectivity of parasite sibships (*z*) and immunity of hosts (*e*) as independent variables to the empirical data in Cornet *et al*. (2009b) (see S3.2, Table S3.1). The fitting was done using the nonlinear least squares (NLS) approach in R (R Core Team, 2024). NLS assumes the residual errors to be normally distributed (Baty *et al*., 2015). We selected the best model based on AICc (corrected Akaike criterion). The fitness function with the least AICc had the form

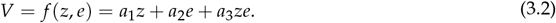

Similar formulations of virulence have been used in prior works (Day *et al*., 2007; Long & Graham, 2011) analysing the combined effects of parasite exploitation of host and host immunopathology on virulence.

To model the effects of niche construction, we replaced the host’s immunity at the individual level *e* in Eq.3.2 with the niche constructed immunity of the same individual host *h*(*z, e*) in the expression for virulence:

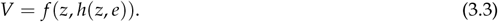

We assumed the niche construction function, *h*(*z, e*), used by parasites (Fig. S3.15) to construct their environment by manipulating the host immunity to be a linear expression of the form

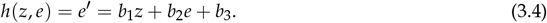

Here, *e*′ is the new immunity of the host post niche construction (or infection), *z* is the corresponding infectivity of the parasite sibship involved in the niche construction and *e* is the immunity of the same host before niche construction (or infection).

For each of eight parasite sibships, Cornet *et al*. (2009b) has data on infectivity of the parasite sibship (*z*), average immunity (after niche construction, *ē*′) of hosts infected by the parasite sibship, and the resulting virulence (see Table S3.1-Supplementary S3.1). This was used to estimate the mean 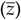 and variance 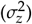 of infectivity. In addition, Cornet *et al*. (2009b) reported the population level mean immunity of the host before niche construction (*e*). This allowed us to estimate the coefficients of the niche construction function *b*_1_, *b*_2_ and *b*_3_ in equation 3.4 (see S3.3, Eq. S3.6). Based on this niche construction function, we could then estimate host immunities before niche construction. We take the variance across these values as environmental variance 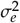 for the sampling approach, but note that it is an underestimation because (Cornet *et al*., 2009a) have averaged over around 100 host individuals.

At the population level, the virulence value can be obtained by non-linear averaging using a Taylor series approximation similar to what is discussed in methods (see sections S3.4 and 2). Since we consider only niche construction in this case study (*g*(*z, e*) = *z*) and the niche construction function considered (Eq. 3.4) is linear for *z* and *e*, we get a reduced form of the expression Eq. 2.3. The expression for average virulence after niche construction is of the form

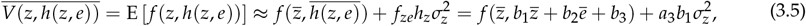

where 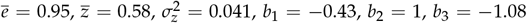 and *a*_3_ = 4.83 (see sections S3.2 and S3.3).

We find that both niche construction and ITV reduce average virulence and thereby increase host and presumably parasite fitness (Fig. 4, Eq 3.5). ITV has no effect on the average virulence in the absence of niche construction. In the presence of both niche construction and ITV, the virulence value the model approximation predicts at population level (≈ 1.004) matches with the value observed empirically (≈ 1.016) (Fig. 4). Also, using our approximation, we compared this with a hypothetical case had the parasite not done niche construction (that is immunodepression upon infection). Had the parasite not done niche construction, the virulence value (= 1.71) would have been higher by ≈ 71% (Fig. 4) compared to the case in which it did niche construction and depressed the immunity of the host. Thus by depressing the immunity via niche construction, the parasite reduced the virulence value (defined as host mortality), in turn increasing its own survival prospectus in the intermediate host (to reach the definitive host to complete its life cycle). Observation of higher virulence in the hypothetical case of parasite not doing niche construction also serves as validation of the hypothesis of immunopathology arrived at in the empirical work (Cornet *et al*., 2009b). Note that because the number of data points in this case study is very small (8), we chose to not perform bootstrapping and thus there are no error bars in figure 4.

**Figure 4.**
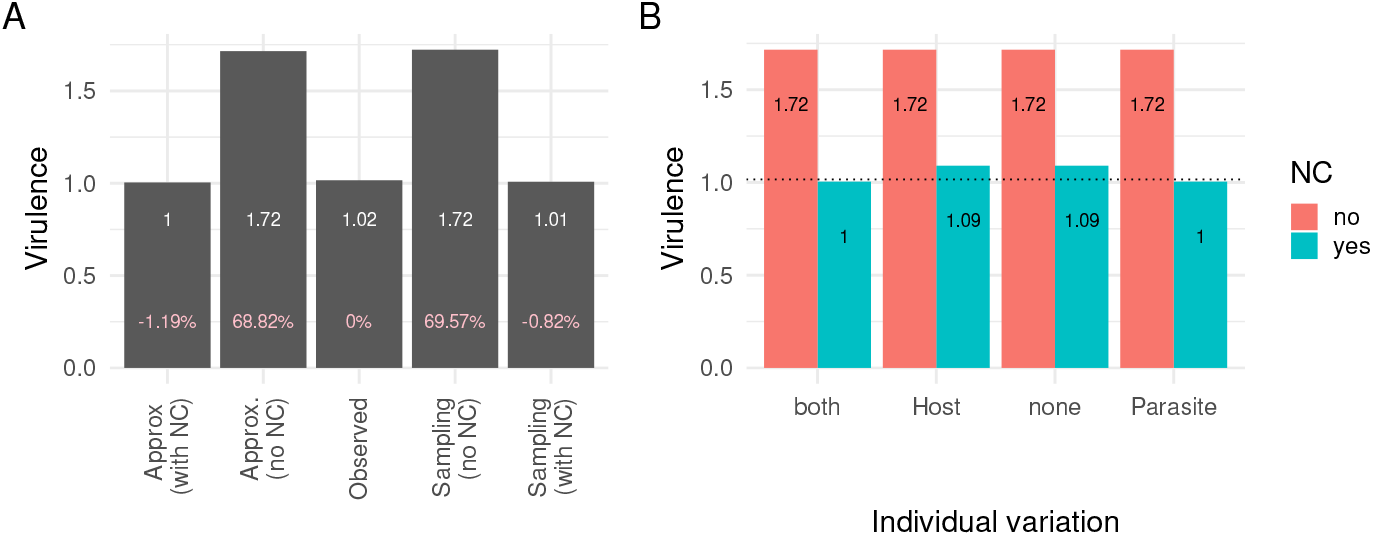
Effect of ITV and Niche Construction: (**A**) Effect of niche construction on average fitness (Virulence). Bars represent (from left to right) the approximation of average fitness with niche construction, approximation of average fitness without niche construction, observed average fitness, simulated average fitness without niche construction, and simulated average fitness with niche construction. ITV is present in both the species. White numbers on the bars are values of average fitness for each bar, pink numbers on the bars are difference between bar value and observed average fitness in percentages of the observed average fitness. (**B**) Effect of ITV and niche construction on average fitness (Virulence) according to the analytical approximation. Bars represent approximated average fitness. Green colour shows the case with niche construction and red shows the case without niche construction. Based on the fitness function and niche construction function considered, host ITV has no effect on virulence. ITV has no effect on virulence in the absence of niche construction. The dashed horizontal line is the observed average fitness.

To study the effects of ITV in virulence, approximations of virulence at population level are computed for cases of no ITV in both species, ITV in either of the two species and ITV in both the species. ITV was found to have an impact on virulence only when niche construction was active (*b*_1_ in Eq. 3.5 is zero without niche construction). Also, ITV in host immunity was found to have no impact on the virulence. This is consistent with Eq 3.5, where only parasite ITV 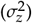 appears. With niche construction active, virulence was found to be least when ITV was present in parasite infectivity and highest in the absence of ITV in parasite infectivity (virulence corresponded to that at mean infectivity of parasites and mean immunity of hosts in population) (Fig. 4).

## 4 Discussion

### 4.1 Discussion of key results

In this paper, we have extended existing non-linear averaging theory for population performance in a variable environment to account for intraspecific trait variation, environmental variation, potentially also among the micro-environments of each individual, niche construction and niche conformance. Using a Taylor approximation or stochastic sampling methods, we can then estimate the effects of ITV, environmental variation, niche construction, and niche conformance on measures of average population performance. The Taylor approximation has many terms due to the high number of processes included, but as seen for the two cases studies, in particular scenarios where only a subset of these processes is relevant, the approximation greatly simplifies and leads then to insightful formulas that allow us to directly see for example whether intraspecific trait variation, niche conformance and niche construction are boosting or reducing average population performance. Thus, the approximation makes it easier to find general principles of how ITV and NC^2^ affect population fitness, compared to, for example, individual-based modelling.

Here, we used two case studies to study the effects of processes operating at the individual level-intraspecific trait variation and niche mechanisms (niche conformance in the first case study and niche construction in the second case study) – on population level demographic parameters (number of fledglings in the first case study and virulence in the second case study) that are shaped by the individual-environment interaction.

In the great tit (*Parus major*) case study, the model estimated the average number of fledglings when there is plastic adjustment of laying dates by birds to match the environment and found the value to be in agreement with the value observed from observed data. We estimated that had the birds not adjusted the laying dates according to the environmental change, the population level fitness (number of fledglings) would have been slightly smaller. The effect of ITV in birds and variation in the environment in the average number of fledglings is dependent on the shape of the fitness function used. Based on the best-fit fitness function we used, both in the presence and absence of niche conformance, ITV in birds had a negative effect on mean number of fledglings. Having some years, when niche conformance has a negative effect on average population fitness (figure S2.12), can be explained by unreliability of environmental cues – a situation when phenotypic plasticity can become maladaptive (Bonamour *et al*., 2019). For this particular population of Great tits it has been shown that environmental cues are quite reliable (Chevin *et al*., 2015), which is in a partial agreement with our results since the fraction of years when niche conformance benefits the population is slightly larger than 50%.

In the host-parasite case study, we studied the *Gammarus pulex* -acanthocephalan (*Pomphorhynchus laevis*) host parasite system that involved a parasite depressing the immunity of its host. To align this with our framework, we perceived the host (immunity as trait) to be the micro-environment for the parasite (infectivity as trait). The individual (parasite)-environment (host) interaction outcome resulted in virulence (host mortality) as fitness outcome. The individual level fitness function was estimated from empirical data. Here we found again that our estimation of the fitness function was in agreement with values observed empirically. In the absence of niche construction by the parasite, the virulence increased, and this difference was more prominent than in the niche conformance case study. In this case, the NC mechanism strongly reduced host mortality by ≈ 70%, which benefits hosts but presumably also parasites. Also, in this case study, we were able to assess the effects of ITV in both species. However, a meaningful effect was achieved only for parasite ITV and only in the presence of niche construction. The direction of the effect on virulence was also negative, as in the niche conformance case study, but in this case this means that parasite ITV improved host survival. In both the case studies, the estimation by the stochastic sampling method was in good agreement to the mean population level values approximated by the Taylor series approximation which shows the accuracy of the approximations (in the range of intraspecific variation and environment variation considered).

How well one can estimate the impacts of niche conformance or construction depends on the type of data available. For example, in the great tit case study, we did not have information on the original traits, i.e. the laying dates before niche conformance. When fitting the niche conformance function, we effectively assumed that an individual’s original trait corresponds to the laying date under the mean environment across the study years. However, in reality, laying dates in the absence of niche conformance might correspond more to the mean environment experienced over an evolutionary time scale and might thus be later. Thus, in reality the birds might perform more niche conformance than we estimate and our estimate of the impact of niche conformance on average fitness should be conservative. In the host-parasite case study, we had at least information on the average environmental value (host immunity) before niche construction, although we did not have the original environmental values of all hosts. Nevertheless, this likely helped us better estimate the impact of niche construction. However, in the host-parasite case study, we did not have immunity trait values for individual hosts, but for averages over around 100 host individuals. Thus the level of host ITV used in the sampling approach is likely an underestimate. However, this problem did not affect the analytical approximation since host trait variance did not appear in the formula. Other empirical studies done so far incorporating ITV in both host and parasite are mostly just considering the genetic diversity of host and parasites (Ganz & Ebert (2010)). In the best case, one would have prior trait values and/or environmental values for all individuals. Of course this is not always possible, especially in field studies, and even in laboratory studies it might require “knocking out” niche conformance or construction in some way.

In the case studies, we have also assumed that there is no evolution within the time scale of the study, which is definitely valid in the host-parasite case study since this involved data from just a single generation. In the case of the great tits, there could in principle be rapid evolution over the multiple-year time span. If some of the changes in laying dates that we see are due to evolution instead of conformance, this could lead to an overestimation of the importance of niche conformance. However, it is not likely in this case since there is no clear trend in green-up date over the duration of the study with plenty of variation. Also, previous studies on great tits have suggested that likely most of the changes in laying date over the past decades are due to plasticity (Charmantier *et al*., 2008).

### 4.2 Future work

Here we have focused on the impact of ITV, niche construction, and niche choice for measures of population performance, such as the average number of fledglings per breeding pair, a measure of reproductive fitness, in the great tit case study and average virulence, a measure of host viability, in the host-parasite case study. Changes in these quantities due to ITV and NC^2^ can then have consequences for population and community dynamics. To model those, future work needs to embed the performance quantity into single- or multi-species ecological models. For the effects of ITV without NC^2^ mechanisms, this has been done e.g. by Hart *et al*. (2016) and van Benthem *et al*. (2024), but it has not been done yet with NC^2^ mechanisms to study e.g. their consequences for population survival or extinction in a changing environment or for species coexistence.

In addition to niche conformance and niche construction, there is a third mechanism by which individuals can achieve a better match between their own trait and their environment within their lifetime: niche choice (Trappes *et al*., 2022), i.e. the selection of an environment. The ability of great tits to change their laying date can be interpreted as temporal niche choice and thus is covered by our model, while including niche choice in a patchy landscape with different environmental conditions in each patch is an open problem for future research. From previous research using other approaches, matching habitat choice is predicted to improve persistence under climate change (Pellerin *et al*., 2019). However, previously optimal habitat choice strategies can also become maladaptive after rapid environmental change (Crowley *et al*., 2019). As mentioned above, assuming a covariance between traits and micro-environments is a simple way of including niche choice into our model. However, as the distribution of micro-environments changes over time, more and more individuals might try to choose the same small number of patches that still have suitable environmental conditions. To understand the consequences of niche choice under environmental change and to avoid overly optimistic predictions, one also has to include competition within patches into the modelling framework.

So far, in our model, niche construction and niche conformance are very similar in their consequences as they both change the distance between an individual’s trait and its micro-environment. When we incorporate patches and modifications of the patch environment also affect other individuals in the same patch, niche construction and niche conformance will then be qualitatively different processes.

Finally, evolution by natural selection is a fourth, though qualitatively different, processes by which the match between individuals and their environment can change (Edelaar & Bolnick, 2019). So far, our modelling approach does not incorporate evolutionary change, but it could be combined with a quantitative genetic model for trait evolution in the future.

## 5 Acknowledgments

We thank Koen van Benthem for advice on statistical analysis, and the members of the Theoretical Biology group of the Bielefeld University, especially Dr. Matthias Spangenberg, for their feedback. This work was funded by the German research foundation (DFG) as part of research consortium SFB-TRR 212, project numbers 316099922 and 396782288.

## Supplementary materials

### S1 Mathematical derivations

#### S1.1 Approximation

The Taylor approximation formula for a function of two variables *f* (*z, e*) around the point (*z, e*) is

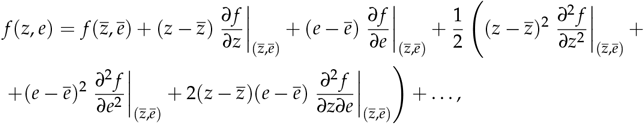

which can be rewritten with different notation as

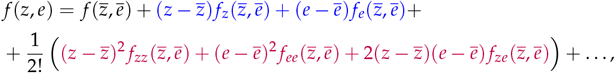

where 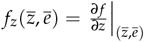, etc.

Now we will do the derivations for the blue and purple parts separately for the case when we have niche conformance and construction functions *g*(*z, e*) and *h*(*z, e*). Suppose *F*(*z, e*) = *f* (*g*(*z, e*), *h*(*z, e*)). Now using the chain rule for total derivates:

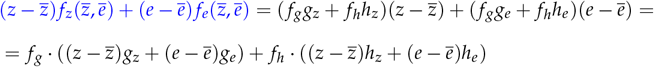

and

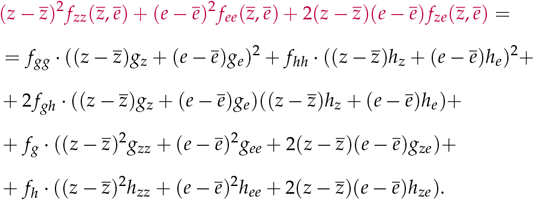

Organising the terms according to derivatives of *f*, we obtain:

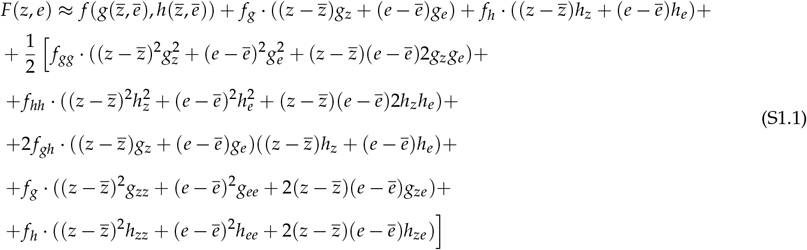

Here by *f*_*gg*_ we mean 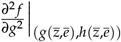, and by *g*_*z*_ we 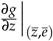 and analogously for *f*_*hh*_, *f* _*gh*_, *g*_*e*_, *g*_*ee*, ∂*z*_ *g*_*zz*_, *g*_*ze*_, *h*_*z*_, *h*_*e*_, *h*_*ee*_, *h*_*zz*_, *h*_*ze*_. Taking the expectation of Eq. S1.1 then gives Eq. 2.3 in the main text.

#### S1.2 Reversing NC2 mechanisms

Suppose we have a study system with NC^2^ mechanisms. For most empirical systems it would be nearly impossible to measure individual trait values without NC^2^ since the observed values are usually the result of these mechanisms. Given, that we measured such trait values we can try to reverse NC^2^ mechanisms to get hypothetical trait values without them.

Let us assume that the trait *Z*′ we observe is in fact a linear combination of some unobserved variable *X* with expectation 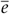 and observable environmental variable *E* (with expectancy *e* and standard deviation *σ*_*e*_): *Z*′ = *X* + *aE*. We have observations of the trait 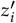 and environment *e*_*i*_ and fit a linear regression 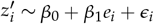 ∼ *β*_0_ + *β*_1_*e*_*i*_ + *ϵ*_*i*_. Then following the principle of the least squares method we minimize the expression 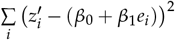 and get the following expressions for the coefficients:

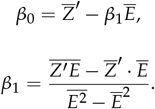

We can find that

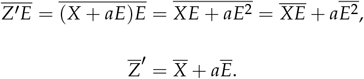

Then

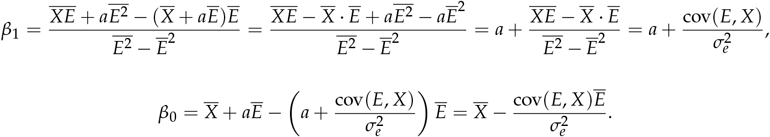

### S2 Niche conformance

#### S2.1 Great tits dataset

We use a dataset published in Cao *et al*. (2019). We go into details on how we cleaned and prepared the dataset in the section Data preparation and provide the prepared dataset with this manuscript.

We choose the following variables from the dataset:

- mother ID,
- year of breeding,
- laying date (as the number of days after March 31, so day 1 = April 1),
- number of fledglings.

For individual-based data we choose only first broods (`Broodtype==0`). Due to the way we model the environment, we take only years of breeding starting with 2001. We use egg laying date as individual trait

We use number of fledglings as fitness proxy.

#### S2.2 Environment dataset

We use NASA satellite green-up data for the years 2001–2021 (Friedl *et al*., 2022). We load the data using the NASA app AppEEARS (AppEEARS Team, 2023). We take coordinates of the study area from (Cao *et al*., 2019), namely, the rectangle between 52°02′ – 52°07′ N and 5°51′ – 5°32′ E. We provide the metadata (file `tits-case-study-MCD12Q2-061-metadata.xml`) and the exact request we used to get the dataset (file `tits-case-study-request.json`) along with the dataset.

#### S2.3 Data preparation

All data cleaning and preparation was performed using the `titsNew data cleaning.R` script.

For the NASA dataset, we worked with the statistics provided with the dataset (file `MCD12Q2-061-Statistics.csv`), since we are interested in yearly average values only. To filter out invalid observations, we removed all rows with less than 40 pixels (`Count` variable). We selected the green-up variable, removing unnecessary columns. Then we converted the date on which the data was collected into the year, and then converted green-up date from days after January 1, 1970 to days after the March 31st for the corresponding year.

**Figure S2.5.**
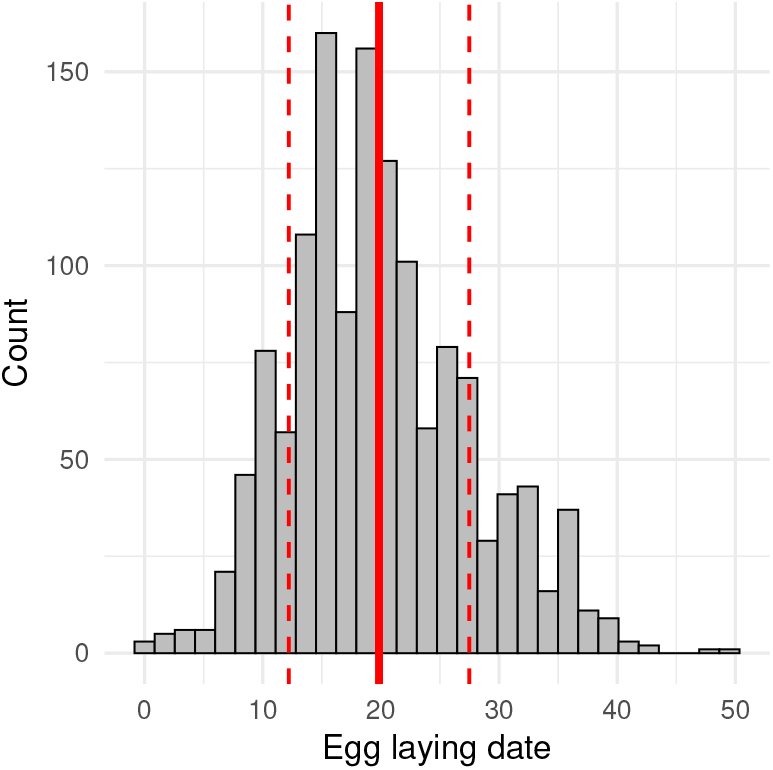
Histogram of the trait (egg laying date in days after March, 31). Trait is on the horizontal axis, count is on the vertical axis. Solid vertical red line is mean trait, dashed vertical red lines are mean plus/minus standard deviation.

**Figure S2.6.**
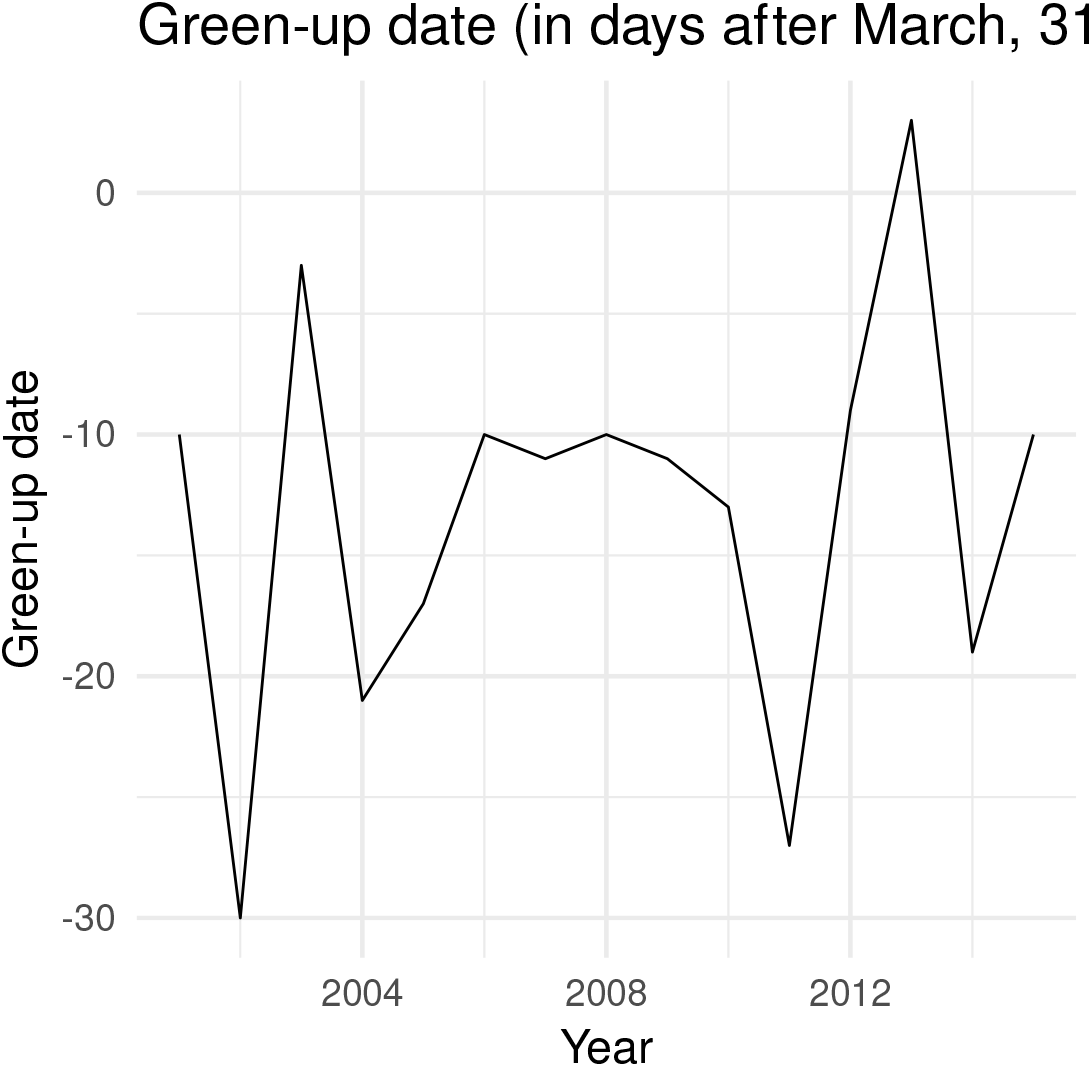
Green-up date (in days after March, 31) for the years 2001–2021. Green-up date is on the vertical axis, year of breeding is on the horizontal axis.

For the great tits dataset, after subsetting the desired variables, we filtered first broods only for years starting from 2001. We also added mother ID as a factor variable. Finally, we added environmental information to this dataset. The laying date data was rescaled by subtracting the mean laying date across the entire data set and dividing by the standard deviation. The green-up date data was analogously rescaled by subtracting its mean and dividing by its standard deviation.

#### S2.4 Model selection procedure

We use generalized linear mixed models (GLMMs) with different distributions (Poisson, zero-inflated Poisson, negative binomial, zero-truncated Poisson) to fit the number of fledglings as dependent variable; laying date (*z*), green-up date (*e*), *z*^2^, *e*^2^, *z* · *e* as fixed factors; and mother ID as random factor. For each distribution we fit all possible fixed parameters combinations with and without the random effect, and for each distribution we choose the model with the best AIC score. Then, among those best models we choose the one that has the best fit in terms of residual plots.

Model selection was done using the `model-selection.R` script. The best model is the zero-inflated Poisson model.

This model consists of two parts: zero-inflated and count. The count part includes intercept, *e, z, z*^2^, *e* · *z*, and the zero-inflated part includes all covariates *e, e*^2^, *z, z*^2^, *e* · *z*, and intercept. We do not have a random effect in this model.

#### S2.5 Estimating the niche conformance function from data

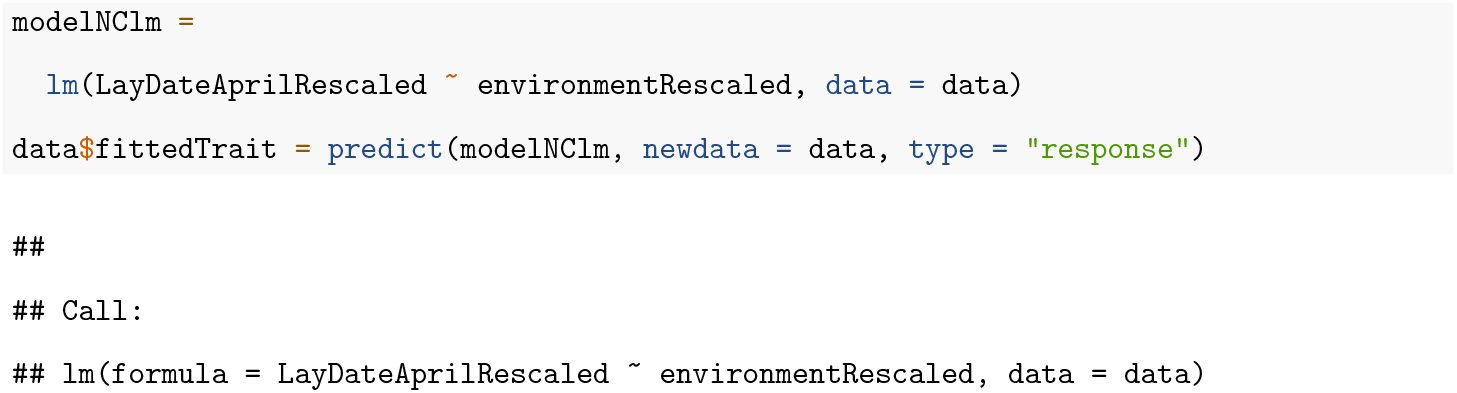

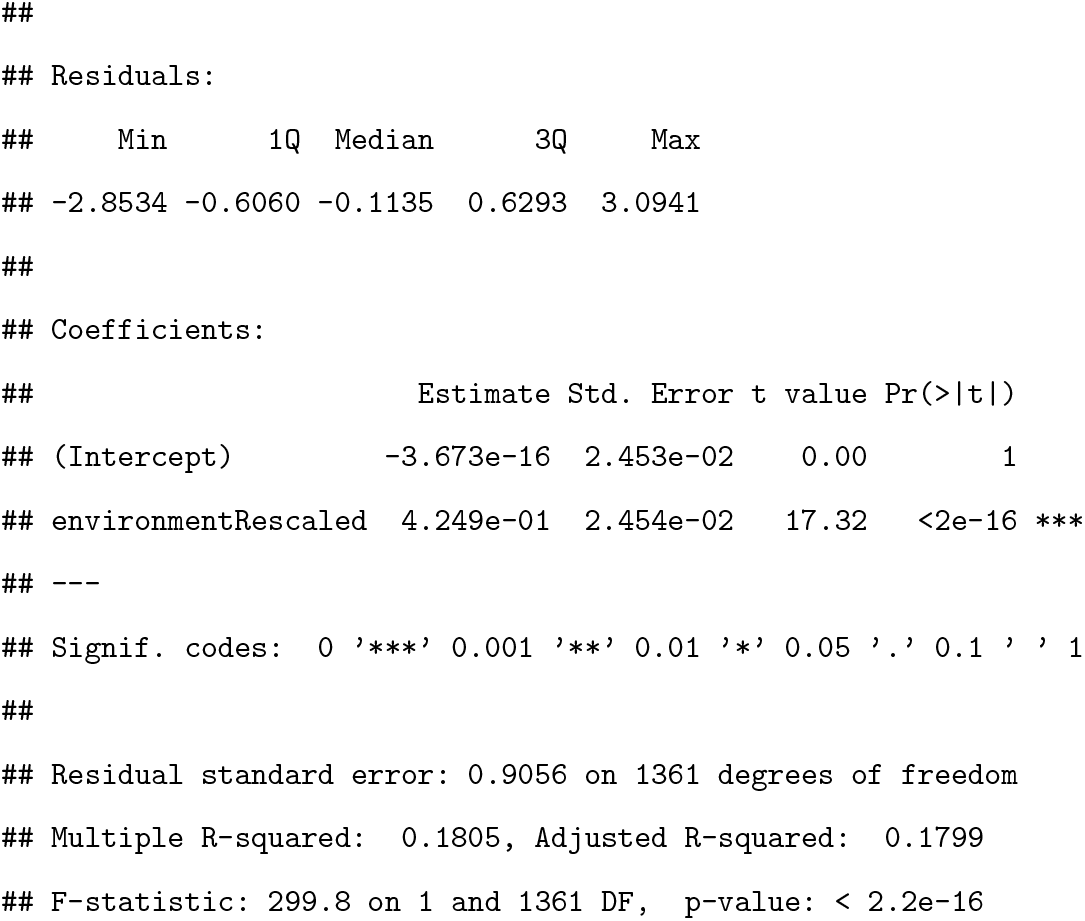

We get the following formula: *z*′ = *a* + *be*. We restore the trait before niche conformance as *z* = *z*′ − *be* (see section S1.2 for more details).

**Figure S2.7.**
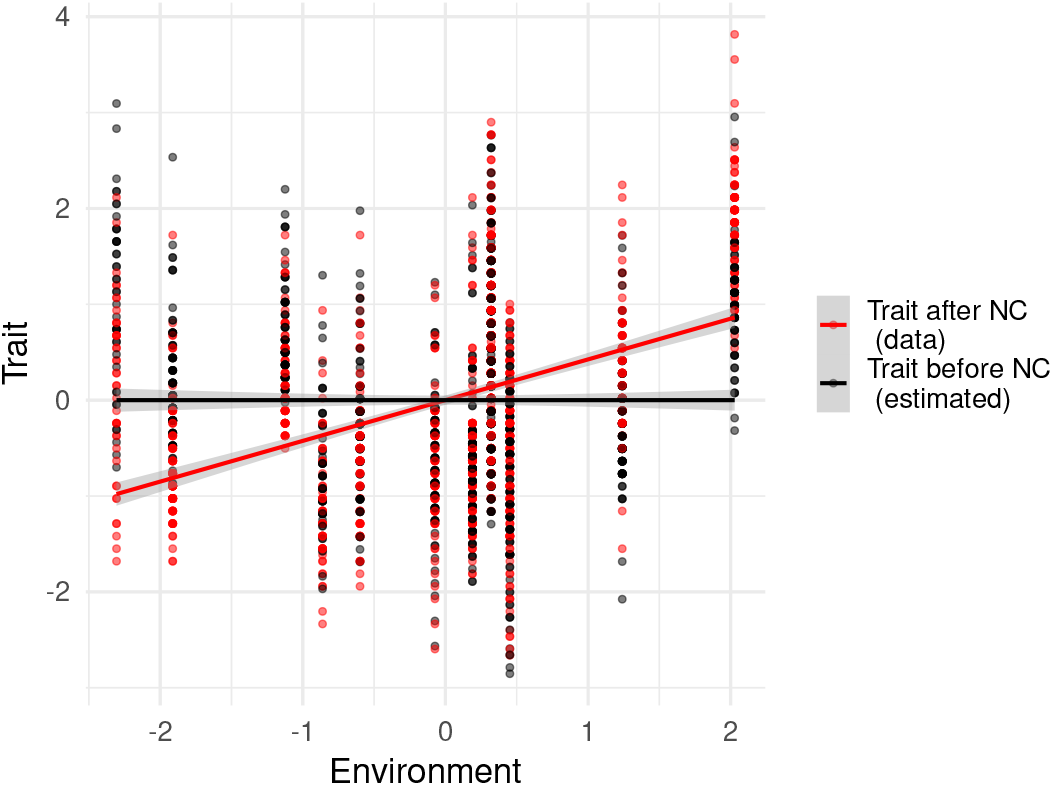
Plot of the trait before (estimated, in black) and after (from data, in red) niche conformance against the environment. Both trait and environment are rescaled.

**Figure S2.8.**
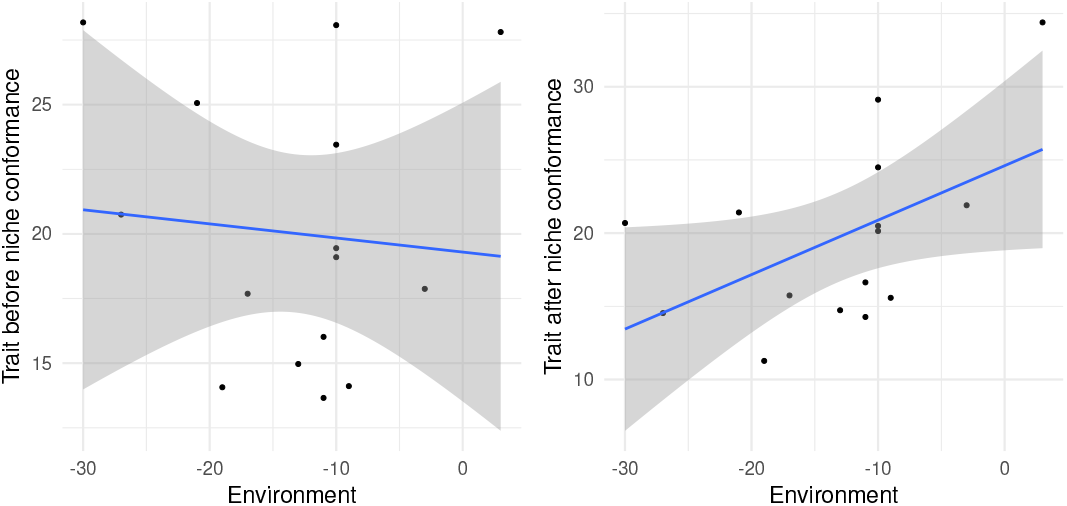
Plot of the average trait before (left) and after (right) niche conformance against the average environment. Each point corresponds to a separate year. The blue line is a simple linear model fit.

#### S2.6 Modelling

We take GLMM’s fixed coefficients to get our fitness function (figure S2.9):

~~~
## exp(1.95115170086993 + -0.079119041927645 * environment + -0.00884878452144596 *
## trait + -0.0398128730304643 * (trait^2) + 0.03201551683258 *
## trait * environment) * (1 - 1/(1 + exp(-(−1.85589924729619 +
## 0.134019589602711 * environment + -0.173064701758474 * (environment^2) +
## 0.038224964584628 * trait + 0.152563049935211 * (trait^2) +
## -0.279784950877135 * environment * trait))))
~~~

To be able to quantify uncertainty in our model estimates, we use a bootstrapping approach. We generate 100 bootstrap samples. For each, we take the prepared dataset and sample rows from it with replacement. The size of the bootstrap sample is the same as the size of the prepared dataset. We refit and save our top GLMM model for individual fitness and the linear model for the niche conformance function for each of the bootstrap samples. We also use the linear model to find trait before niche conformance (also for each bootstrap sample). We do all of that in the `titsNew bootstrap samples.R` script.

We then check the bootstrap convergence by calculating average fitness via approximation on bootstrap samples of different size going from 1 to 100 (`titsNew bootstrap convergence.R`). We do that by calculating the approximation for every bootstrap sample with fitness and niche conformance functions estimated from that sample. We provide average laying date as the new trait value for our approximation function, trait before niche conformance as the old trait, and average green-up date as both old and new environment value.

**Figure S2.9.**
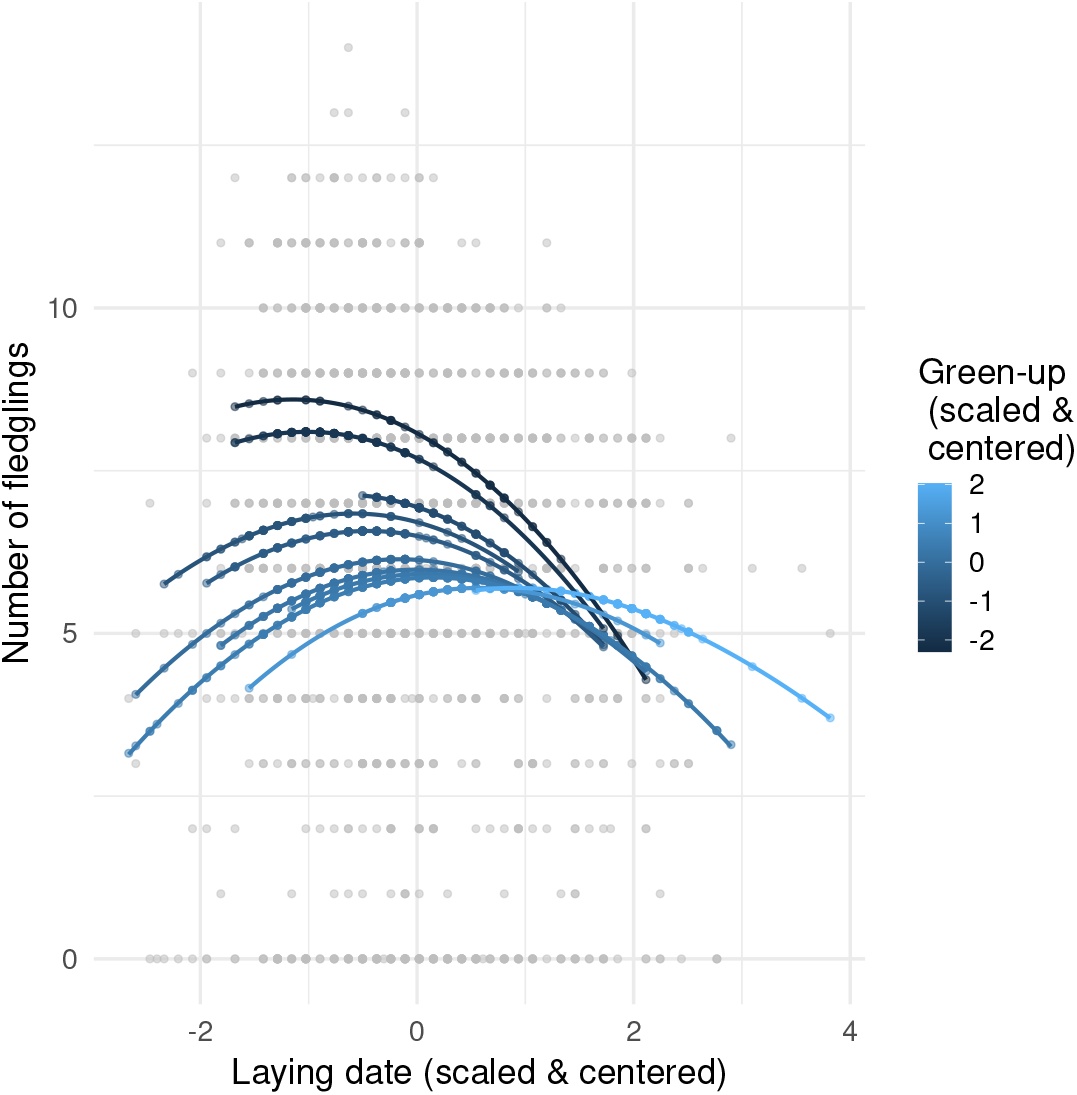
Plot of the 2D fitness function. In a very light grey are the actual data points. Points in shades of blue are fitted numbers of fledglings. Blue lines are the result of geom smooth applied to fitted values.

**Figure S2.10.**
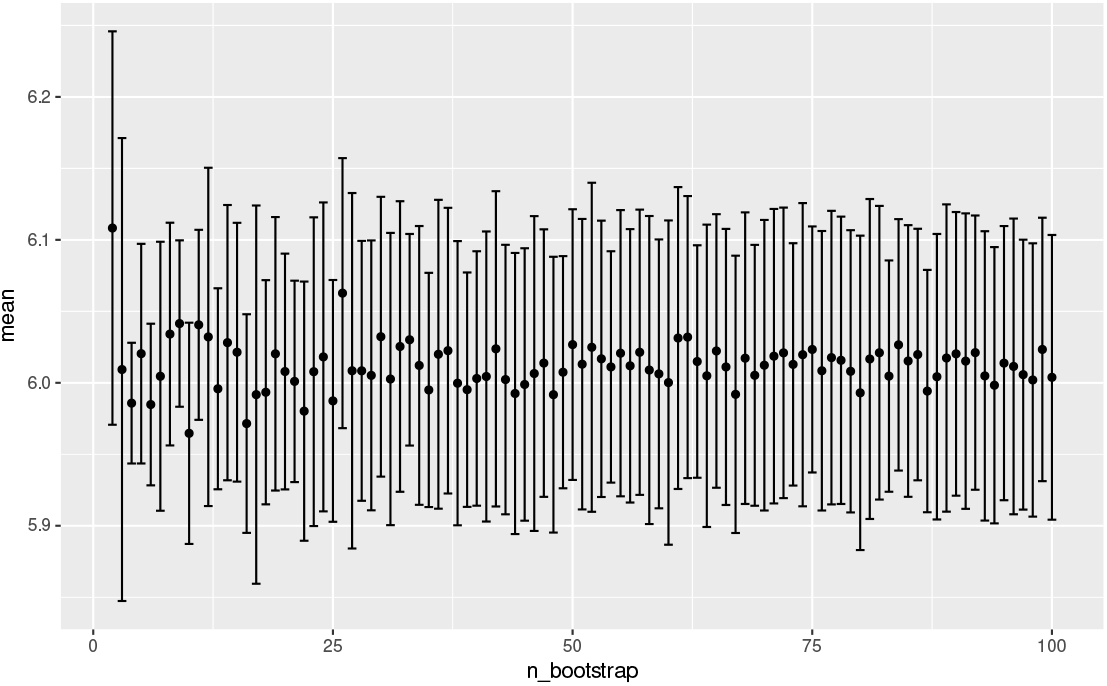
Convergence of the approximation method. The horizontal axis is the number of bootstrap samples used to calculate the approximation, the vertical axis is the average number of fledglings, the error bars are plus/minus standard deviation.

#### S2.7 Modelling for each year

The code used to produce the figures in this subsection is in `titsNew bootstrap hist1.R` for figures S2.11 and 3A and `titsNew bootstrap hist2.R` for S2.12 and 3B.

First, we compare how well the analytical approximation works compared to the simulation method, and how both methods estimate fitness in a hypothetical case of absence of niche conformance. To have an idea of how sensitive the approximation and the simulation method are, we perform bootstrapping.

We use the sampling approach to provide a baseline to which we can compare how well our approximation method performs. For that, we randomly draw individual traits and calculate their fitness for the given environment in each year (function `simulationYearlyFunction()` in the `titsNew functions.R` script).

We use the same number of individuals and the same environment value as in the original dataset. We assume that individual traits are normally distributed with mean and standard deviation calculated from the dataset narrowed down to the corresponding year, using the traits before niche conformance. We start our simulation procedure by drawing individual traits. Then we calculate traits after niche conformance. If we aim to include niche conformance in the simulation, then we use the niche conformance function estimated from the whole dataset. Otherwise, we use the expression *z*′ = *z* meaning that the trait does not change. After that, we calculate individual fitness by simply feeding the environment vector and the vector of traits after niche conformance into the fitness function estimated from the whole dataset. The output of the simulation function is one number: average fitness across individuals.

**Figure S2.11.**
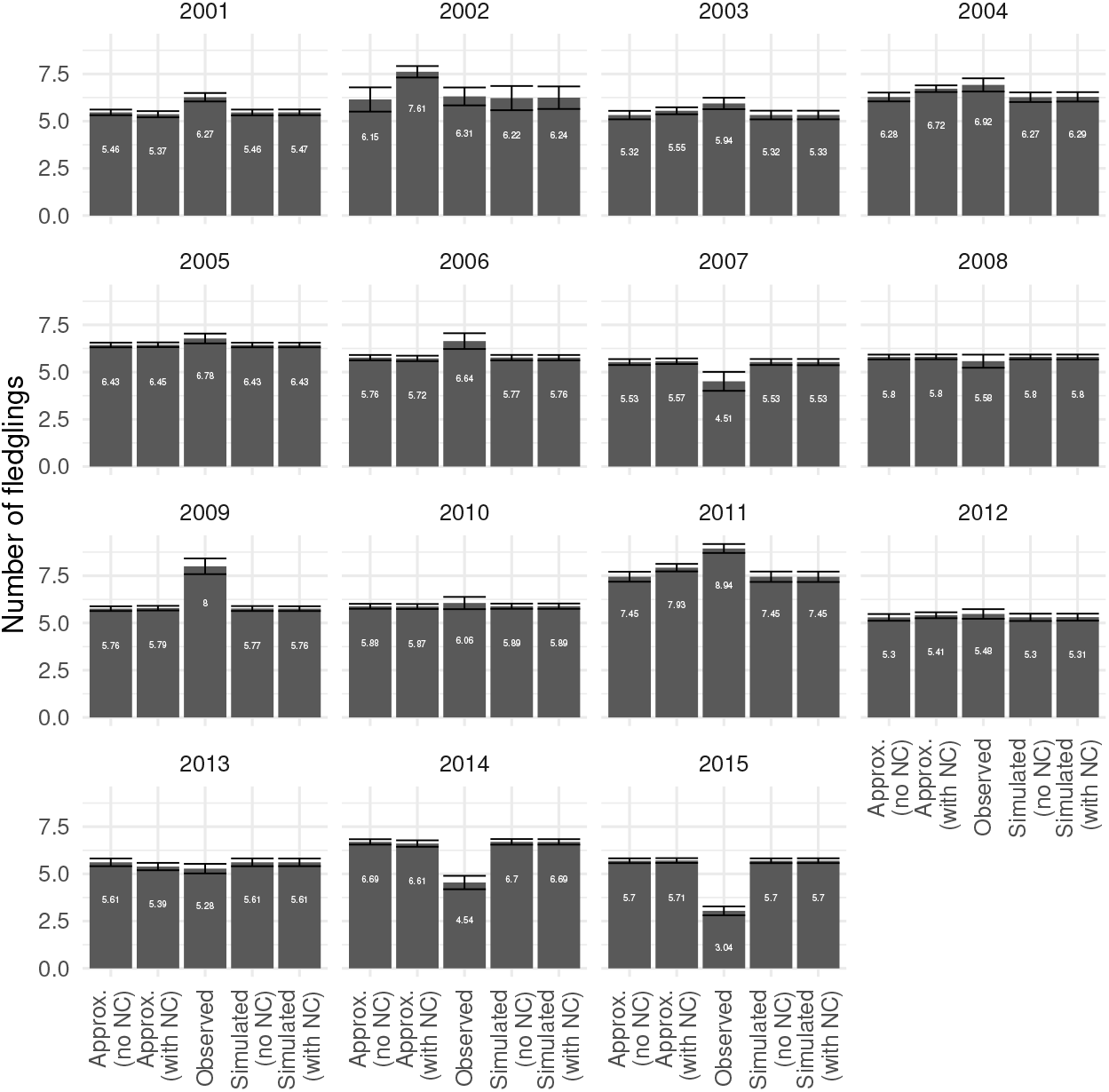
Histogram similar to figure 3A, but calculated for each year separately, shows effects of niche conformance on average fitness. On each facet bars represent (from left to right) approximation of average fitness without niche conformance, approximation of average fitness with niche conformance, observed average fitness, simulated average fitness without niche conformance, simulated average fitness with niche conformance. White numbers on the bars are values of average fitness for each bar. Error bars represent uncertainty due to bootstrapping procedure as standard deviation (for approximation and simulated average) and standard error of the mean (for observed average).

**Figure S2.12.**
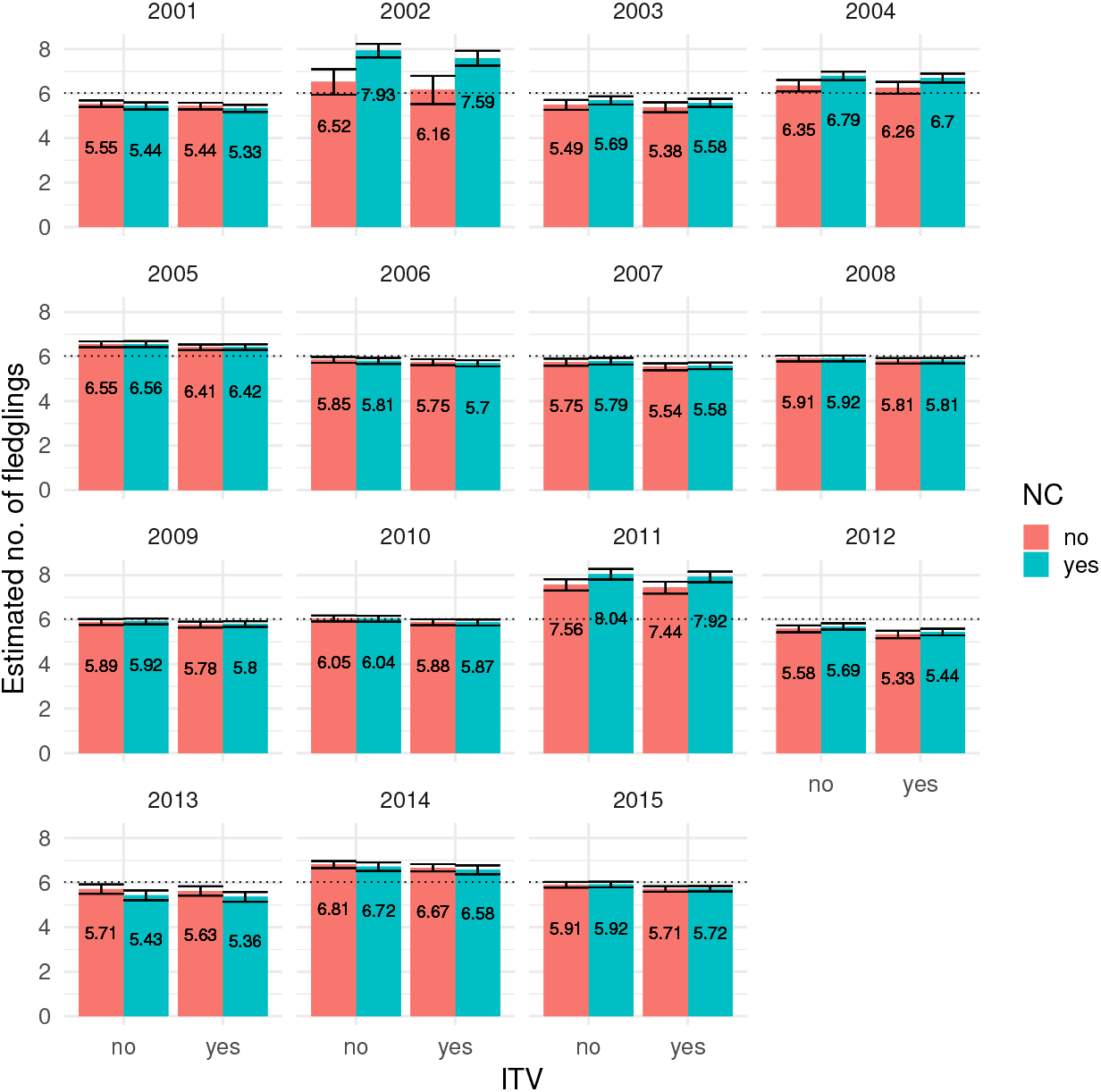
Histogram similar to figure 3B, but calculated for each year separately. In each facet the effect of ITV and niche conformance on individual fitness according to analytical approximation is shown. Bars represent approximated average fitness. The two leftmost bars in each facet represent the case without ITV, the two rightmost bars represent the case with ITV. The two red bars represent the case without niche conformance, the two green bars represent the case with niche conformance. Numbers on the bars are values of approximated average fitness for each bar. Error bars represent uncertainty due to bootstrapping procedure as standard deviation. The dashed horizontal line is the observed average fitness

We get values for analytical approximation of average number of fledglings for each year, and for each method (approximation and simulation) we get the standard deviation across bootstrap samples.

#### S2.8 Modelling for the whole dataset

We do the same analysis as for the previous section, not for each year separately, but for the whole dataset (this is the analysis underlying Fig. 2 in the main text).

We use again the simulation method in almost exactly the same fashion as for yearly simulations. We use the same sampling approach, by randomly drawing individual traits and calculating their fitness for the given environment at each year, but now averaging the fitness over all 15 years. To do that we use the function `simulationAllFunction()` from the `titsNew functions.R` script.

We use the same overall number of individuals and the same environment values as in the original dataset. We assume that individual traits are normally distributed with mean and standard deviation calculated from the dataset using traits before niche conformance. For each year, we draw the number of individuals for this year from the Poisson distribution with mean equal to average number of individuals in a year from the dataset. We start our simulation by generating the sample of individuals that will be used for our simulations (not every individual will appear in every year). We start our simulation procedure by drawing individual traits for the whole sample of our individuals, and randomly sample a number of them determined by the Poisson distribution. Then we calculate traits after niche conformance. If we aim to include niche conformance in the simulation, then we use the niche conformance function estimated from the dataset. Otherwise, we use expression *z*′ = *z* meaning that the trait does not change. After that we calculate individual fitness by simply feeding the environment vector and the vector of traits after niche conformance into the fitness function estimated from the dataset. The output of the simulation function is one number: average fitness for all 15 years.

### S3 Niche construction

#### S3.1 Extraction of data from Cornet et al. (2009b)

Immunity of the host, infectivity of the parasite sibship infecting that particular host and the corresponding virulence is obtained from Cornet *et al*. (2009b). In Cornet *et al*. (2009b), we use the data of only the spring experiment. Figure 2 in Cornet *et al*. (2009b) gives the infectivity of each parasite sibship. Figure 3b in Cornet *et al*. (2009b) gives the average immune proxy (proPO activity) values of hosts infected by the parasite sibships. Figure 4 in Cornet *et al*. (2009b) gives the virulence value corresponding to the immune values (proPO activity) of the hosts. Get Data Graph Digitizer software (Informer Technologies, 2015) was used to get the data points from Cornet *et al*. (2009b). A collated information of the immunity of the host, infectivity of the parasite sibship and resulting virulence sourced from Cornet *et al*. (2009b) is given in Table S3.1.

#### S3.2 Fitting and model selection

To estimate the interaction function, we fitted several polynomial interaction functions to virulence (*V*), immunity of host (*e*) and infectivity of the parasite sibship (*z*) data in Table S3.1 using nonlinear least squares (NLS) in R. The models compared were

**Table S3.1.** Data collated from Cornet *et al*. (2009b): For each parasite sibship, the immunity of the hosts infected by parasites of that sibship, infectivity of the parasite sibship, and the resulting virulence as taken from Cornet *et al*. (2009b). Immunity is measured as natural logarithm of proPO activity. Infectivity is defined as the number of infected hosts divided by the total number of surviving hosts at the end of the experiment. Virulence is defined as the parasite-induced host mortality (risk ratio).

| parasite sibship | immunity (post niche construction) | infectivity | virulence |
| --- | --- | --- | --- |
| E | -0.755 | 0.81 | 0.68 |
| D | -0.51 | 0.83 | 0.75 |
| A | -0.37 | 0.22 | 0.96 |
| C | -0.57 | 0.45 | 1.01 |
| B | -0.27 | 0.49 | 1.04 |
| F | -0.041 | 0.53 | 1.1 |
| G | -0.38 | 0.71 | 1.27 |
| H | -0.15 | 0.60 | 1.32 |

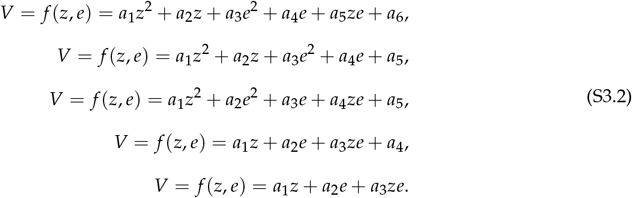

Model selection was performed using AICc (the corrected Akaike criterion) in R. The selected model, i.e. the one with the least AICc, was

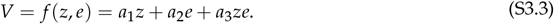

with parameter values *a*_1_ = 2.18, *a*_2_ = −2.33, *a*_3_ = 4.83 (Fig. S3.13).

#### S3.3 Construction function

The immunity *e* of the host is assumed to be niche-constructed (depressed) by the parasite sibship with infectivity *z* according to the function

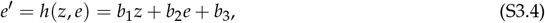

where, *e*′ is the new immunity of the host after niche construction and *b*_1_, *b*_2_ and *b*_3_ are constants. Had the immune values of the individual hosts before and after niche construction be known, the constants could have been estimated by fitting Eq. S3.4 to data sets of *e*′, *e* and *z*. However, Cornet *et al*. (2009b) report only the mean immunity value *ē* = 0.95 of all the hosts prior to niche construction and average immunities across approximately 100 host individuals (*ē*′) after niche construction, as well as the infectivities (*z*) of the respective individual parasite sibships (Table S3.1). To estimate the immunities of the host individuals before niche construction (*e*), we fitted a linear regression of the form

**Figure S3.13.**
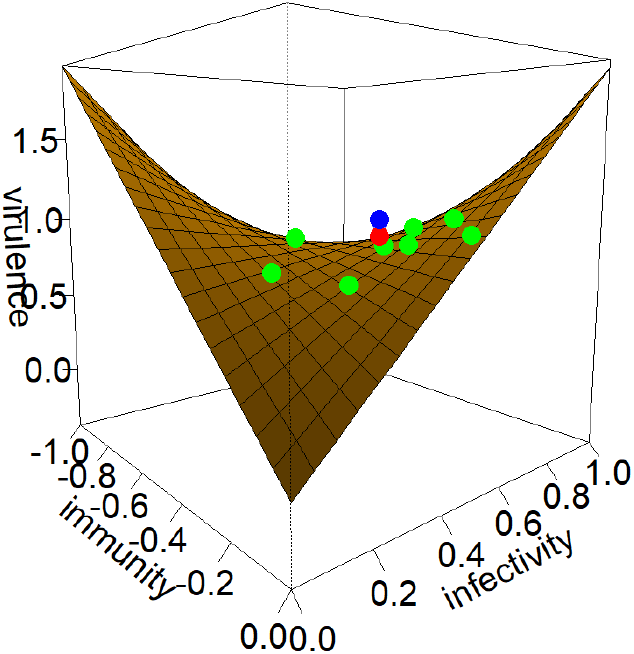
Fitness function: Function (Eq.S3.3) fitted to the immunity of host individuals, infectivity of the corresponding parasite sibships and the resulting virulence. Green points are the individuals. The red point stands for the virulence value (0.91) corresponding to the mean immunity and mean infectivity. The blue point stands for the mean virulence value (1.01) observed empirically. The values differ because of the effect of non-linearity in interaction (Jensen’s inequality)-The interaction outcome corresponding to the mean traits across individuals is not equal to the mean of individual interaction outcomes.

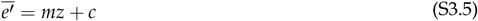

to the data for average immunity after niche construction (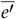) and infectivity (*z*) (Table S3.1, blue line in figure S3.14). Estimated values of slope and intercept were *m* = −0.43, *c* = −0.129. In addition, we assume that deviations of individual host immunities from the mean host immunity before niche construction, *e* − *ē*, are preserved after niche construction. Thus, we obtain

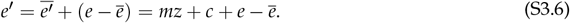

**Figure S3.14.**
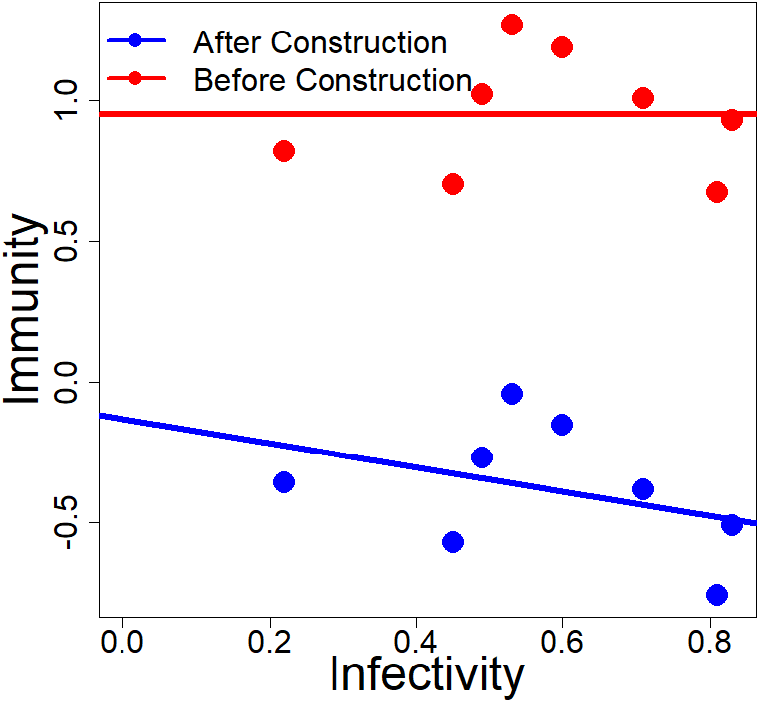
Estimation of the niche construction function: The blue dots are the immunity values of the host from the data in Cornet *et al*. (2009b) (see Table S3.1). We estimated the immunity before niche construction which are shown as red dots. The red solid line corresponds to the average immunity value of the host prior to niche construction (*e*=0.95) and the solid blue line corresponds to the best fit linear regression between average immunity after niche construction (*ē*′) and infectivity.

Comparing Eq. S3.6 and Eq. S3.4 we get

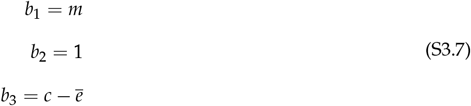

Estimated values of the constants were *b*_1_ = −0.43, *b*_2_ = 1, *b*_3_ = −1.08 (figure S3.15).

Now, based on Eq. S3.6, we could estimate the immune values of hosts before niche construction as *e* = *e*′ − *mz* − *c* + *e* (see Fig. S3.14) and Table S3.1 could be modified to include it as Table S3.2.

#### S3.4 Approximation

We apply the general framework as given by Eq. 2.4 to get the non-linearly averaged fitness function at the population level. In the context of the niche construction case study, this will give the expression for average virulence. Since niche conformance is not considered in this case study, *g*(*z, e*) = *z*. Thus, we get *g*_*e*_=0 and *g*_*z*_=1. Also, since we use a niche construction function which is linear with respect to the variables *z* and *e*, many terms in Eq. 2.3 become 0 and hence the expression simplifies. When the fitness function is given by Eq. 3.2 and the niche construction function is given by Eq. 3.4, we will get *f*_*zz*_=0, *f*_*ee*_=0, *g*_*z*_=1, *g*_*e*_=0, *g*_*zz*_=0, *g*_*ee*_=0, *g*_*ze*_ = 0, *h*_*zz*_=0, *h*_*ee*_=0 and *h*_*ze*_=0. We take *Cov*(*z, e*)=0 as host immunity prior to niche construction and parasite sibship infectivity are independent entities and parasites are assumed to have no choice in infecting hosts with particular immunity. The Taylor series approximation of virulence thus simplifies to,

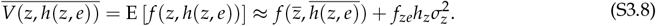

**Table S3.2.** Collated data from Cornet *et al*. (2009b) with estimated host immunities before niche construction: For each parasite sibship, the average immunity of the hosts infected by the parasite sibship, the infectivity of the parasite sibship, and the resulting virulence as taken from Cornet *et al*. (2009b). Estimated immunities of the host prior to infection are also included. Immunity is measured as the natural logarithm of proPO activity. Infectivity is defined as the number of infected hosts divided by the total number of surviving hosts at the end of the experiment. Virulence is defined as the parasite-induced host mortality.

| immunity, $e$ (pre niche construction) | immunity, $e'$ (post niche construction) | infectivity, $z$ | virulence, $V$ |
| --- | --- | --- | --- |
| 0.6738258 | -0.755 | 0.81 | 0.68 |
| 0.9266411 | -0.51 | 0.83 | 0.75 |
| 0.8179316 | -0.37 | 0.22 | 0.96 |
| 0.7027883 | -0.57 | 0.45 | 1.01 |
| 1.0204573 | -0.27 | 0.49 | 1.04 |
| 1.2663271 | -0.041 | 0.53 | 1.1 |
| 1.0044961 | -0.38 | 0.71 | 1.27 |
| 1.1875327 | -0.15 | 0.60 | 1.32 |
| Mean $\bar{e}=0.95$ | | Mean $\bar{z}=0.58$ | |
| Variance $\sigma_e^2=0.045$ | | Variance $\sigma_z^2=0.041$ | |

Substituting the values of 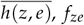 and *h*_*z*_ we get

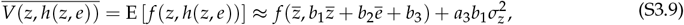

where *b*_1_ = −0.43, *b*_2_= 1, *b*_3_ = −1.08 and *a*_3_ = 4.83. 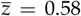 and 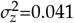 are computed from the infectivity column in table S3.1 and *e* = 0.95 directly from Cornet *et al*. (2009b).

**Figure S3.15.**
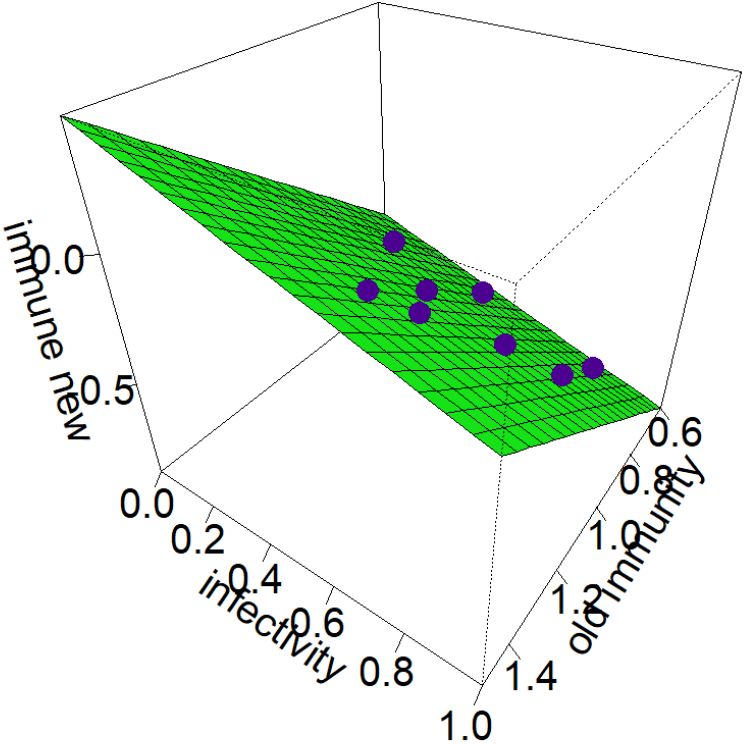
Niche construction function: At the individual level, niche construction is done by the parasite sibship with infectivity (*z*) on a host of immunity (*e*) to change the host immunity from *e* to *e*′. Each of the blue points gives relationship between average host immunity before and after niche construction when infected with a parasite of specific infectivity.

## Notes

### Competing Interest Statement

The authors have declared no competing interest.

